# Extending eco-evolutionary theory with oligomorphic dynamics

**DOI:** 10.1101/2022.12.12.520082

**Authors:** Sébastien Lion, Akira Sasaki, Mike Boots

## Abstract

Understanding the interplay between ecological processes and the evolutionary dynamics of quantitative traits in natural systems remains a major challenge. Two main theoretical frameworks are used to address this question, adaptive dynamics and quantitative genetics, both of which have strengths and limitations and are often used by distinct research communities to address different questions. In order to make progress, new theoretical developments are needed that integrate these approaches and strengthen the link to empirical data. Here, we discuss a novel theoretical framework that bridges the gap between quantitative genetics and adaptive dynamics approaches. “Oligomorphic dynamics” can be used to analyse eco-evolutionary dynamics across different time scales and extends quantitative genetics theory to account for multimodal trait distributions, the dynamical nature of genetic variance, the potential for disruptive selection due to ecological feed-backs, and the non-normal or skewed trait distributions encountered in nature. Oligomorphic dynamics explicitly takes into account the effect of environmental feedback, such as frequency- and density-dependent selection, on the dynamics of multi-modal trait distributions and we argue it has the potential to facilitate a much tighter integration between eco-evolutionary theory and empirical data.

## 1 Introduction

A key challenge in deepening our understanding of the evolution of quantitative traits in nature is the development of theory that captures the key role of ecological processes on evolutionary change and integrates this with data. Both natural selection and genetic drift are fundamentally ecological processes driven by the population dynamics of genetically diverse populations. Furthermore, gene flow is often determined by ecological processes such as dispersal. Moreover, by altering the genetic makeup of populations, micro-evolutionary processes in turn change the ecological conditions, which leads to a feedback loop between ecology and evolution. There is a long history of studying these feedbacks (Pimentel, 1961; Hutchinson, 1965), leading to classic work on density- and frequency-dependent selection (Chitty, 1967; Clarke, 1972), and to the development of theoretical frameworks aiming to capture eco-evolutionary feedbacks (for recent reviews, see McPeek (2017), Lion (2018), Govaert et al. (2019), and Klausmeier et al. (2020)). The two prominent approaches for studying the evolution of quantitative traits in an ecological context are quantitative genetics (QG) and adaptive dynamics (AD).

Both QG and AD are versatile frameworks and can be applied to a variety of ecological or demographic models, allowing for discrete- or continuous-time dynamics, discrete population structure (as in matrix models, Caswell (2001)) or continuous population structure (as in integral projection models, Rees & Ellner (2016)). As noted by several authors, these two frameworks generate similar long-term predictions under the assumptions of small mutation effects (for AD) and narrow trait distributions (for QG) (Abrams et al., 1993; Abrams, 2001; Day, 2005; Lion, 2018). However, AD and QG models have often been developed by distinct research communities, originally with different types of questions in mind. For instance, QG was initially developed to study the short-term transient dynamics resulting from standing genetic variation, and the changes in mean trait of a population over a few generations (Lande, 1976, 1979; Walsh & Lynch, 2018). A large part of this literature has considered simple ecological scenarios with simple forms of density-dependence, often ignoring frequency-dependence (Box 1), as in classical moving-optimum QG models (Kopp & Matuszewski, 2014). In contrast, adaptive dynamics is tailored to investigate long-term eco-evolutionary endpoints under mutation-limited evolution, using the notion of ‘invasion fitness’ as its main fitness concept (Metz et al., 1992; Geritz et al., 1998; Dercole & Rinaldi, 2008; Metz, 2011).

The recent surge of interest in rapid evolutionary processes (Thompson, 1998; Hairston et al., 2005; Kopp & Matuszewski, 2014; Hendry, 2017; Govaert et al., 2019; Bassar et al., 2021) has led many researchers to use frequency-dependent QG models to jointly consider the ecological and evolutionary dynamics of a focal population (see e.g. Mougi & Iwasa (2010), Schreiber et al. (2011), Vasseur et al. (2011), Cortez & Weitz (2014), Patel & Schreiber (2015), Klauschies et al. (2016), Cortez (2018), Yamamichi et al. (2019), and van Velzen et al. (2022)). Indeed, AD always assumes that evolution is relatively slow with ecological dynamics reaching equilibrium before the potential invasion of a new mutation and therefore cannot examine the impact of rapid evolutionary change. However, these eco-evolutionary QG models assume a narrow unimodal trait distribution with constant variance, and approximate the eco-evolutionary dynamics through the joint change of population density and trait mean, neglecting the effect of selection on the shape of the distribution. As a result, they cannot capture the emergence of multimodal distributions through frequency-dependent disruptive selection, or changes in the skewness of the distribution. Empirical evidence of skewed (Bonamour et al., 2017) or multimodal distributions (Emlen, 1994; Duffy et al., 2008; Anderson et al., 2009) in the wild suggests that the widely used assumptions of Gaussian or narrow unimodal distributions with constant variance may be too restrictive. On the other hand while AD is able to capture the diversifying processes leading to polymorphism through the phenomenon of evolutionary branching, it does so in the limit of vanishingly small standing variation.

There is therefore the need for an integrative framework to model and understand eco-evolutionary dynamics (Box 1) across different time scales (e.g. fast vs. slow evolution) and to take into account the effect of directional, stabilising, and disruptive selection on realistic distributions of quantitative traits. Various authors have attempted to build bridges between AD and QG (Abrams et al., 1993; Abrams, 2001; Day, 2005; Kremer & Klausmeier, 2013; Cortez, 2016; Lion, 2018), but they have typically done so by highlighting the connections between these two approaches and not by suggesting a new framework to go beyond the current limitations of both theories. Our goal in this perspective is to discuss a theoretical development which aims to bridge the gap between QG and AD approaches. We do this in the simpler case of clonal reproduction (a widely used assumption in AD) because, although the classical QG approach considers sexually reproducing organisms, most eco-evolutionary QG approaches do not explicitly model the effect of sexual reproduction on population dynamics and yield equations which are identical to those derived for asexual organisms. This simplification will also make the approach directly applicable to a wide range of classical ecological models.

The approach we describe here uses an “oligomorphic” approximation introduced by Sasaki & Dieckmann (2011), and is related to various trait-based approaches developed in community ecology and quantitative genetics (Wirtz & Eckhardt, 1996; Norberg et al., 2001; Débarre et al., 2013, 2014; Barabás & D’Andrea, 2016; Débarre & Otto, 2016; Klauschies et al., 2018; Mullon & Lehmann, 2019; Barabás et al., 2022; Wickman et al., 2022). This new theoretical framework explicitly takes into account the effect of environmental feedback, notably frequency- and density-dependent selection, on the joint dynamics of ecological variables and trait distributions, and has the potential to facilitate a tighter integration between eco-evolutionary theory and empirical data. Here, we show how this framework can be used to analyse the eco-evolutionary dynamics across different time scales, as it allows us to examine both fast evolutionary dynamics in non-equilibrium population (typically analysed in ecological and quantitative genetics models) and slow evolutionary dynamics in populations that have reached an ecological attractor (typically analysed using adaptive dynamics). We also discuss how this approach can be used to extend quantitative genetics theory to account for the dynamical nature of genetic variance, the potential for disruptive selection due to ecological feedbacks, and the non-normal or skewed trait distributions typically encountered in nature. We conclude by discussing the links with other established theoretical methods and by highlighting some persepectives for further theoretical developments and applications.

**Box 1: Glossary**

**Eco-evolutionary feedback**: Stricto sensu, this refers to the reciprocal interaction between ecological and evolutionary processes.

**Eco-evolutionary dynamics**: We use this term to describe the coupled dynamics of ecological variables and of genetic or phenotypic distributions, due to eco-evolutionary feedbacks. Although some authors have advocated a narrow use of the term, restricted to cases where there is no separation in time between ecological and evolutionary dynamics (e.g. rapid evolution; Bassar et al. (2021)), our definition is broader as we wish to emphasise the continuum between dynamics with a pure separation of time scales and dynamics with overlapping time scales, and the ability of OMD to describe how variation in the relative time scales of ecological and evolutionary proceses can lead to different forms of eco-evolutionary feedback.

**Environmental (or ecological) feedback:** Fitness is a property of a geno- or phenotype (with trait value *z*) measured in a given environment **E**, hence the notation *r*(*z*, **E**) we use in this manuscript. The environment **E** is affected by the actions of the individuals and by the composition of the population (densities, frequencies of types, trait distribution), and this in turn feeds back on fitness. The notion of environmental feedback thus generalises the concepts of density-dependence or frequency-dependence.

**Frequency-dependence**: Frequency-dependence is an ubiquituous but ambigous concept (Heino et al., 1998). Stricto sensu, it means that fitness depends on the frequencies of the different genotypes in the population. In realistic ecological scenarios with environmental feedback, this is always the case, simply because the fitness of a given individual is affected by the presence of other individuals with different traits. For instance, in an AD setting, a mutant that has a positive invasion fitness at low frequency will have fitness zero after fixation. Frequency-dependent QG models typically assume that the environmental feedback can be summarised by the mean trait 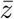 and therefore use a fitness function of the form 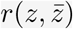, which is a special case of the general environmental feedback formulation. For our purpose here, it will be sufficient to equate frequency-independence with the absence of environmental feedback (i.e. fitness can be written as a function of the phenotype only, *r*(*z*), but see Metz & Geritz (2016) for a more refined definition).

## 2 Classical eco-evolutionary approaches

In this section, we will briefly review the three classical approaches to model eco-evolutionary dynamics, which are population genetics, adaptive dynamics, and quantitative genetics. Readers interested in a more thorough discussion of the differences and similarities between these theoretical frameworks may look up Abrams (2001), Day (2005), and Lion (2018). For didactic purposes, we will focus on simple versions of these theories. For instance, as typical of many eco-evolutionary models, we will consider asexual populations. We will also ignore much of the complexities caused by genetic architecture, population structure, or the effects of the environment on the genotype-phenotype map. We refer the reader to section 5 for a more detailed discussion of these issues.

**Box 2: Two models of resource competition**

**Model 1: adaptation of a resource exploitation trait** As our main running example, we use a classical model of consumers exploiting a single dynamic resource, with density *R*. We assume that the consumer population is genetically diverse, and we consider the following ecological dynamics for the density *n_i_* of consumers with phenotype *z_i_*:

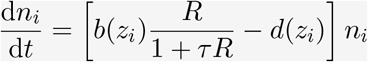

where *b*(*z*) and *d*(*z*) are the fecundity and death rates of the consumer population, which depend on the phenotype *z*, and the link between resource and consumer reproduction follows a type-II functional response with attack rate 1 and handling time *τ*. We couple the consumer dynamics to an equation describing the production and consumption of the resource (see Online Appendix S). In Model 1, we investigate the evolution of reproductive effort, that is we consider a trait *z* that affects both fecundity *b*(*z*) or mortality *d*(*z*) and ask whether the consumer should preferentially invest into reproduction or survival, given the ecological context it experiences.

**Model 2: evolutionary diversification through niche partitioning** To investigate frequency-dependent disruptive selection, we use the classical MacArthur-Levins model of resource competition, analysed using OMD by Sasaki & Dieckmann (2011). The consumer dynamics are given by

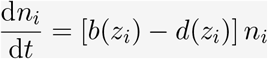

where the phenotype *z* is a resource utilisation trait. Individuals with similar phenotypes exploit similar resources (or niches) and thus compete more intensely than individuals with distant phenotypes. This is captured by an increased mortality rate. For instance, with a discrete phenotypic distribution, we write

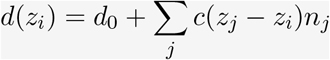

where *d*_0_ is the baseline mortality rate, and *c*(*z_i_* − *z_i_*) is a function that quantifies the strength of competition experienced by a type-*i* consumer from a type-*j* consumer, and is assumed to depend on the difference between the phenotypes of the two types. Assuming a trade-off between competitive ability and fecundity, Model 2 can lead to diversification with different morphs occupying different niches.

We will use a classical resource-consumer model (Model 1 in Box 2) as a running example, but our main argument will use the following general formulation

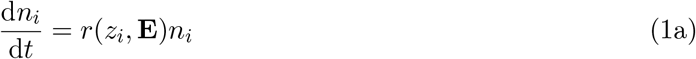

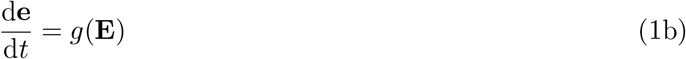

where *n_i_* is the density and *r*(*z_i_*, **E**) the per-capita growth rate of individuals with phenotype *z_i_*. The function *r*(*z*, **E**), for a given phenotype *z* experiencing environment **E**, will play an important role and will often be called “fitness function” in the rest of this article. As in Lion (2018), we follow the typical practice in adaptive dynamics of making the fitness function depend on the trait and on the environment, **E**, thereby materialising the environmental feedback (Box 1). The vector **E** contains the densities *n_i_* and any extrinsic environmental variable necessary to calculate fitness, collected in a variable **e** (for instance, in Model 1 in Box 2, **e** is simply the density of resource *R*).

The function *g* captures the dynamics of **e** and will in general depend on all the phenotypes in the population, although we leave this dependency implicit for simplicity. For completeness, we note that the environmental feedback formulation generalises the practice of writing fitness as a function of the individual phenotype *z* and the mean phenotype 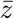 in frequency-dependent QG models (Box 1; see also e.g. Iwasa et al. (1991), Day (2005), and Lion (2018)).

The formulation in equation 1 should be simply understoood as a general framework for eco-evolutionary change that describes how the densities of individuals with particular phenotypes change depending on their per-capita growth rates, which are functions of the environment **E**. The critical point is that the environment varies due to in part to feedbacks between ecological and evolutionary processes. Starting with this general formalism, the different analytical methods (population genetics, quantitative genetics, and adaptive dynamics) typically make particular assumptions that we will describe in detail below.

### 2.1 Population genetics

Our first approach will be to track the change in the frequencies of the different types, thereby following the central idea of classical population genetics (PG). Defining *n* = ∑_*i*_ *n_i_* as the total density of consumers, and *f_i_* = *n_i_*/*n* as the frequency of type *i*, we obtain

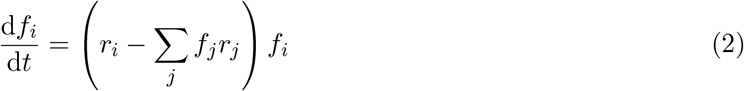

where *r_i_* = *r*(*z_i_*, **E**) is a short-hand for the per-capita growth rate of type *i*. The second term between brackets is the average growth rate of the total population, 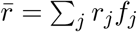. For our running example, we have

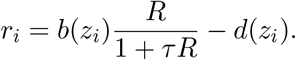

Equation (2) is a version of the replicator equation (Taylor & Jonker, 1978) and simply tells us that type *i* will increase in frequency if its per-capita growth rate is greater than the average growth rate of the population. Note that in this version of the replicator equation, the change in frequency depends on the environmental feedback (e.g. through the density of resources *R*). In the special case where we only have two types (a wild-type *w* and a mutant *m*), the change in mutant frequency is simply given by

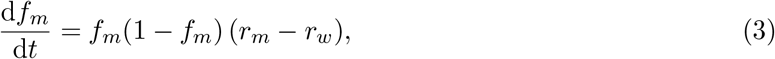

where *f_m_*(1 − *f*_m_) measures the genetic variance in the population, and *r_m_* − *r_w_* is the selection coefficient. This is a classical result of population genetics, which can be obtained either in discrete or in continous time, but note that equation (3) does not assume that the selection coefficient is a constant and takes into account ecological feedbacks (through the density of resources in our example). Thus, equations (2) and (3) need to be coupled to dynamical equations for the ecological densities (e.g. the total density of consumers *n* and the density of resources *R*). This approach has been used to study the interplay of ecology and single-locus genetics (see e.g. (Charlesworth, 1971; Roughgarden, 1971; Yoshida et al., 2007; Yamamichi et al., 2011; Yamamichi, 2022)).

### 2.2 Adaptive dynamics

The key assumption of adaptive dynamics (AD) is that mutations are rare. This leads to a separation of time scales where mutants arise at low frequency in a resident population on its ecological attractor. Hence, evolution is mutation-limited and unfolds on a much slower time scale than ecological processes. This assumption is justified by limiting arguments on the mutation process (Metz et al., 1992; Geritz et al., 1998; Rousset, 2004; Lehmann & Rousset, 2014; Van Cleve, 2015) and has fostered the development of a detailed toolbox to deal with evolution with complex ecological interactions.

In adaptive dynamics, the success of a mutant allele is measured by its invasion fitness (Metz et al., 1992; Dieckmann & Law, 1996; Metz et al., 1996; Geritz et al., 1998), which is explicitly defined as a function of the ecological variables characterising the resident community. Hence, invasion fitness clearly captures the notion of environmental feedback (Metz et al., 1992; Mylius & Diekmann, 1995; Ferrière & Legendre, 2013; Lion, 2018), although it does so under the assumption of mutation-limited evolution.

In our example, it is straightforward to calculate the invasion fitness from equation (3). Assuming that the resident attractor is an equilibrium, we have *r_w_* = 0 and therefore the mutant increases in frequency if *r_m_* is positive. Invasion fitness can thus be written in terms of the mutant trait *z_m_* and of the eequilibrium resident environment 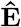 as

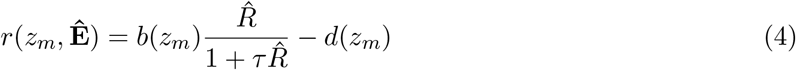

Because the density of resources at equilibrium 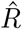 is a function of the resident trait *z_w_*, it is also often convenient in practice to define invasion fitness as a function of the traits, as *s*(*z_m_*, *z_w_*).

It is beyond the scope of this perspective piece to give a full overview of the AD toolbox, and we will instead focus on two key results that will be important in the following. Both results are derived under the additional assumption that mutations have small phenotypic effects (often called weak selection in the literature, Rousset, 2004), so that *z_m_* − *z_w_* is small. The first result is that, under this assumption, the direction of selection is given by the selection gradient, which is the derivative of invasion fitness with respect to the mutant trait, evaluated at neutrality (i.e. when the mutant and resident traits are equal). The zeros of the selection gradient then correspond to evolutionary singularities. The second result is that the evolutionarily stability of a singularity can be assessed through the sign of the second derivative of invasion fitness.

For our example, the selection gradient is

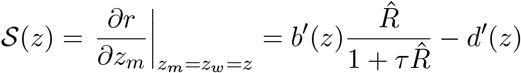

and depends on the slopes of fecundity and mortality with respect to the phenotype, while the second derivative evaluated at a point *z*^⋆^ such that 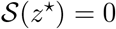 is

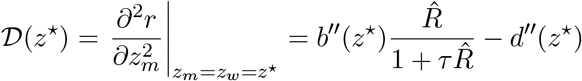

and depends on the curvature of the life-history trait functions.

Adaptive dynamics has become a widely used approach to study long-term phenotypic evolution in the presence of eco-evolutionary feedbacks. However, there are key limitations of adaptive dynamics as a model of eco-evolutionary dynamics that is of use to empiricists. First, it is not well suited to study short-term evolutionary dynamics fueled by standing genetic variation in the population. Second, theoretical investigations of evolution in polymorphic resident populations, although conceptually well established (Geritz et al., 1998; Kisdi, 1999; Durinx et al., 2008), are often mathematically complex and restricted to potential evolutionary endpoints. Third, invasion fitness, as a theoretical construct, is difficult to integrate with empirical or experimental data. In practice, in most systems, it is often difficult to carry out a large number of reciprocal invasion experiments between pairs of different variants, as we typically rarely have multiple phenotypes characterised, or good conditions under which invasion can be quantified, and, even if we do, the scale and duration of the experiments would often be impractical.

### 2.3 Quantitative genetics

Originally, quantitative genetics (QG) was specifically developed to model short-term genetic and phenotypic changes in natural populations. The aim of QG is to track the dynamics of the distribution of a quantitative trait determined by many loci with small phenotypic effects. Instead of considering a discrete distribution of types (through the frequencies *f_i_*’s), we now assume a continuous distribution *ϕ*(*z, t*), normalised such that ∫ *ϕ*(*z, t*)d*z* = 1 over the trait space, and model how this distribution changes over time. Often, this is a very difficult task and the majority of studies focus on the dynamics of moments (characteristics) of this distribution, typically the mean trait or (less frequently) the variance of the trait (Bulmer, 1971; Lande, 1976, 1979; Lande & Arnold, 1983; Barton & Turelli, 1987; Bürger, 1991; Walsh & Lynch, 2018).

In QG, the dynamics of the mean trait are given by the Robertson-Price equation (Robertson, 1966; Price, 1970; Queller, 2017; Lion, 2018) which relates the change in the mean trait 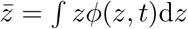 to the covariance between the trait and the per-capita growth rate of a given type, and from which the well-known breeder’s equation can be derived (Walsh & Lynch, 2018). We have

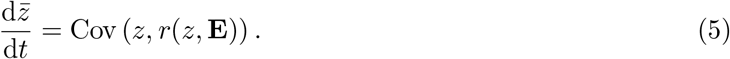

The dependency on the environmental feedback **E** shows that QG models explicitly handle the coupling between ecology and evolution. For instance, in our resource-consumer example, we can couple the dynamics of the mean trait with that of the total density of the population (Slatkin, 1980; Taper & Case, 1992; Day, 2005). However, in contrast to AD, this coupling does not, without additional assumptions, rely on a separation of time scales, which allows for the study of short-term evolution resulting from existing genetic variation in the population.

In order to make progress however, most QG models make additional assumptions, in the wake of Lande (1976, 1979, 1982)’ seminal work. One central and very common assumption is that the trait distribution is and remains normally distributed (Lynch & Walsh, 1998; Walsh & Lynch, 2018). This allows tractability and leads to considerable insight. However, it also puts major constraints on the evolutionary process, and prevents QG models from examining how multimodal distributions can be generated by frequency-dependent disruptive selection, or how selection acts on skewed or multi-modal trait distributions. Another classical assumption of QG models is that selection is frequency-independent. This yields dynamical equations for the mean and higher moments of an arbitrary trait distribution for weak (Barton & Turelli, 1987; Turelli & Barton, 1990; Bürger, 1991) or strong (Turelli & Barton, 1994) selection. However, with frequency-independent selection, only specific forms of ecological feedbacks can be analysed.

In order to study eco-evolutionary processes, researchers have attempted to partly relax these assumptions and turned to a frequency-dependent version of QG which assumes that the trait distribution is tightly clustered around its mean (Iwasa et al., 1991; Abrams et al., 1993; Vincent et al., 1993; Taylor & Day, 1997; Day, 2005). This small variance approximation leads to the following approximation of equation (5)

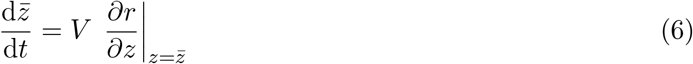

where *V* is the (additive) genetic variance. Equation (6) shows that the mean trait changes along the slope of an adaptive landscape, but note that the selection gradient is given by the slope of the individual fitness function instead of the mean population fitness (Iwasa et al., 1991; Abrams et al., 1993; Day, 2005). For our running example, this leads to

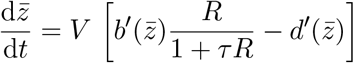

The direction of selection is thus given by a selection gradient which takes the same form as in AD, with two distinct features. First, the density of resources *R* is not necessarily at equilibrium. Second, the rate of evolution is scaled by the existing standing variation through the genetic variance *V*. (Note that, here, heritability is assumed to be 1 for simplicity, so that there is not distinction between genetic and phenotypic variance, but extensions are discussed in section 5.1).

Many QG models take a further step and consider that the variance is constant (but see e.g. Bulmer, 1971; Lande & Arnold, 1983; Barton & Turelli, 1987; Bürger, 1991; Turelli & Barton, 1994; Walsh & Lynch, 2018). Clearly, such approximations will fail once the distribution of a quantitative character becomes bimodal as is expected under frequency-dependent disruptive selection, but also when one is interested in understanding how selection alters the shape of trait distributions. The need for an alternative approach is highlighted by empirical evidence of skewed (Bonamour et al., 2017) or multimodal distributions (Emlen, 1994; Duffy et al., 2008; Anderson et al., 2009).

### 2.4 The need for an integrated approach

What is currently the best tool to study the interplay of ecological and evolutionary processes? The answer clearly depends on the biological question we want to solve. Ecological models or PG single-locus models have been used to explore rapid evolutionary dynamics due to ecologically mediated competition between genotypes, but they are less suitable to study the longer-term dynamics of quantitative traits fuelled by standing variation or recurrent mutations. AD provides a solid mathematical framework to study long-term evolution in the presence of environmental feedbacks, but does so using the assumption that evolution is limited by rare mutations, and is therefore not well suited to make predictions on short-term evolution, as typically observed in the field or in experiments. On the other hand, QG is perfectly suited to investigate rapid evolutionary dynamics in complex ecological scenarios, but in practice, most models rely on some restrictive assumptions. For instance, a large body of literature focuses on models of adaptation to a constant or moving optimum (for a review, see Kopp & Matuszewski (2014)). In such models, the optimum is externally determined by the abiotic environment and the fitness function is Gaussian or quadratic. Such a priori constraints prevent the investigation of more complex feedback loops between the abiotic or biotic environmental factors and the evolution of quantitative traits. On the other hand, frequency-dependent QG models can be used to explore more complex eco-evolutionary dynamics (e.g. Patel & Schreiber (2015)), but these models typically assume narrow unimodal trait distributions with constant variance, and therefore cannot handle the dynamics of multimodal or non-Gaussian distributions, or the emergence of polymorphism under frequency-dependent disruptive selection.

Thus, we think there is value in trying to bridge the gap between AD and QG approaches. As we have just seen, bridging this gap is facilitated by the fact that, in eco-evolutionary theory, AD and QG use similar assumptions (e.g. weak selection) and share key concepts (e.g. the selection gradient). The weak selection assumption stems from either assuming that mutation effects are small or that standing genetic variation is small. As a result, both approaches capture the effect of directional selection through a gradient formulation: the change in the mean trait is given by a measure of genetical variation multiplied by a selection gradient which gives a first-order (linear) approximation of fitness (Abrams, 2001; Day, 2005; Lion, 2018). However, assuming weak selection affects the time scales between ecological and evolutionary dynamics, and therefore the shape of environmental feedbacks. This is because evolution by natural selection will be slower when the amount of genetic variation is small. One may therefore wonder whether this assumption, despite its ubiquity and technical usefulness, is really suited to study rapid evolutionary dynamics, often caused by high standing variation and/or mutation rates. In the next section, we show how a small variance assumption can be coupled with the oligomorphic approximation introduced by Sasaki & Dieckmann (2011) to help us move beyond both the focus on unimodal character distributions with constant variance, typically encountered in QG, and on mutation-limited evolution, typical of AD.

## 3 Multi-morph eco-evolutionary dynamics

The key idea underpinning oligomorphic dynamics (OMD) (Sasaki & Dieckmann, 2011) is to decompose a multimodal trait distribution into a sum of narrow unimodal morph distribution. Intuitively, each morph corresponds to one peak of the trait distribution, and is characterised by its abundance (e.g. its frequency), its position (e.g. its mean trait value), and its width (e.g. its standard deviation). By deriving the dynamics of the morph frequencies, mean trait values, and variances, it is thus possible to determine how each peak changes or moves over time, and therefore how the full trait distribution, at the population level, changes as well.

To explain these ideas, we will give a brief overview of the general framework, before showing how it can be applied to shed light on various aspects of eco-evolutionary dynamics.

### 3.1 The oligomorphic decomposition

The first step is to decompose a possibly multimodal trait distribution into several morphs. A morph is a cluster of continuous variants around a phenotypic mean trait. Mathematically, we write

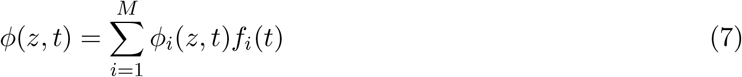

where *ϕ*(*z, t*) is the trait distribution at the population level, *M* is the number of morphs, *ϕ_i_*(*z, t*) is the trait distribution of morph *i*, and *f_i_*(*t*) is the frequency of morph *i* (figure 1a). A frequent simplifying assumption will be that the morph distributions *ϕ_i_*(*z, t*) are Gaussian, but this is not required for most of the theory.

**Figure 1:**
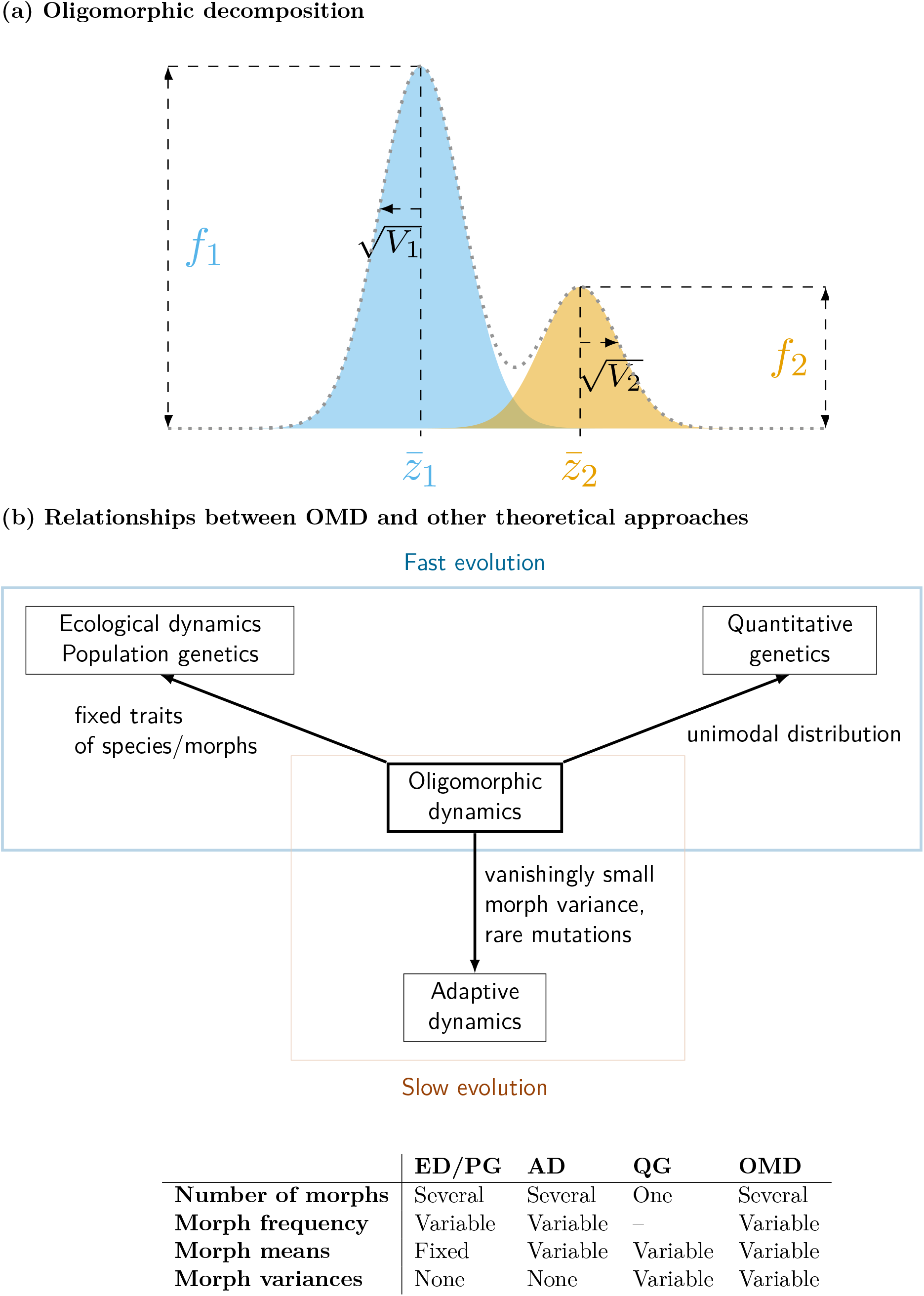
OMD in a nutshell. (a) The full trait distribution *ϕ*(*z, t*) (dotted line) is decomposed into a sum of the morph distributions *ϕ*_1_(*z, t*) and *ϕ*_2_(*z, t*) (with means 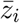 and variances *V_i_*), weighted by the morph frequencies *f*_1_ and *f*_2_. The blue and orange shaded regions correspond to *f*_1_*ϕ*_1_(*z, t*) and *f*_2_*ϕ*_2_(*z, t*) respectively. (b) Relationships between OMD and other theoretical approaches.

An intuitive interpretation of the decomposition (7) is that the mean value of a morph should correspond to one of the modes of the distribution. However, the oligomorphic decomposition does not require that the distance between the morphs is large, and is also valid when the population contains two very similar morph distributions with different frequencies. Hence, the decomposition allows for a quasi-monomorphic trait distribution, with two nearly identical morphs, to split into two modes through disruptive selection. The required number of morphs will depend on the biological scenario (see section 5.4).

### 3.2 The small morph variance approximation

Once our trait distribution is formally split into several morphs, we further assume that each morph is tightly clustered around its mean, 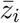, so that the quantities 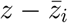 are all proportional to a small parameter *ε*. This is equivalent to saying that the morph variances, *V_i_*, are of order *ε*^2^. With this assumption, we can apply a Taylor approximation to the per-capita growth rate *r*(*z*, **E**) around the mean of each morph, and use this approximation to derive the dynamics of the ecological densities, morph frequencies, and morph moments. A short overview is given in Appendix A and we focus on the key results below.

### 3.3 Dynamics of ecological densities

Equipped with this approximation, we can now derive the dynamics of the total population density, *n*(*t*). We obtain

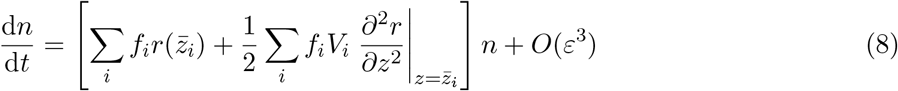

Here, as in all equations from now on, we drop the explicit dependency of fitness on **E**, and only write *r*(*z*) to simplify the notation.

To leading order, the per-capita growth rate of the total population density can thus be approximated by the sum of the growth rates at the morph means, 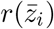, weighted by the morph frequencies, *f_i_* (the first term between brackets). For fixed morph means, this exactly corresponds to the dynamics of the total density of a multi-species population, as encountered in community ecology.

The second term between brackets is scaled by the morph variances and therefore corresponds to a second-order correction. It captures the decrease in the mean growth rate 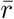 due to individual phenotypic deviations from the mean trait, known as the “genetic (or standing) load” in the QG literature (Lande & Shannon, 1996; Kirkpatrick & Barton, 1997; Chevin, 2013; Wickman et al., 2022). In equation (8), the genetic load is a multi-morph extension of these previous results and takes the form of an average, over all morphs, of the strength of stabilising or disruptive selection around the morph mean (given by the curvature of the fitness function *r*(*z*)), weighted by the morph genetic variance. In practice, this second-order term can often be neglected with good accuracy, as in the simulations used below, but it would be interesting to investigate the conditions for which taking into account the impact of genetic variation on population dynamics becomes important (Bolnick et al., 2011).

### 3.4 Dynamics of morph frequencies

Equation (8) depends on the dynamics of morph frequencies and morph means. The dynamics of morph frequencies take the following form

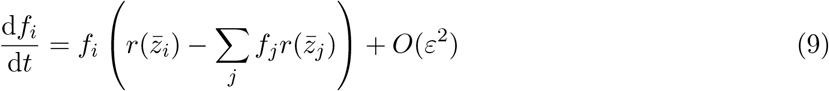

To leading order, this equation takes exactly the same form as the equation governing allele frequency change in population genetics (e.g. equation (2)). However, in the oligomorphic approximation, the morph means are not fixed, so the changes in morph frequencies depend on how the peaks of the multimodal distribution move over time. Competitive exclusion of one morph corresponds to a decrease to zero of that morph’s frequency.

### 3.5 Dynamics of morph means

The next step is therefore to derive equations for the dynamics of the morph means. Using the small morph variance approximation, we obtain

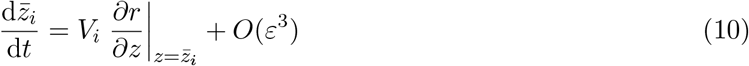

Thus, to leading order, the rate of change in morph means is scaled by the amount of genetic variation, measured by the morph variance *V_i_*, and its direction is given by the selection gradient (the slope of the fitness function *r*(*z*) evaluated at the morph mean). We thus recover a morph-specific version of equation (6), with the key difference that the selection gradient may also depend on the means of the other morphs, so that the equations for the morph means are all coupled. Equation (10) tells us in which direction the modes of our distribution move. The long-term endpoints are given by the zeros of the selection gradient, and correspond to the predictions of an AD analysis. As shown by Sasaki & Dieckmann (2011), the condition for the stability of the dynamics of the morph means is equivalent to the convergence stability condition of AD.

### 3.6 Dynamics of morph variances

In contrast to the typical practice in QG, the oligomorphic approach does not assume that the morph variances are fixed. Hence, we can go one step further and derive the following equation for the change in morph variances, with the additional assumption that morph distributions are all symmetric around their means:

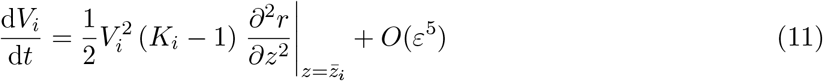

where *K_i_* is the kurtosis of the morph distribution. Thus, the rate at which the genetic variance of a morph changes is scaled by the amount of genetic variation *V_i_*, and by the degree of presence of outliers in the trait distribution, measured by the kurtosis *K_i_* (so that the change in variance will be faster if the distribution has more outliers). The direction of the change in the genetic variance of the morph is given by the sign of the curvature of the fitness function *r*(*z*) at the morph mean. As shown by Sasaki & Dieckmann (2011), the condition for the stabililty of morph variances corresponds to the evolutionarily stability condition of AD. Note that, in discrete time, the dynamics of variance have an extra term corresponding to the square of the selection gradient (see Online Appendix S for a discussion).

### 3.7 Moment closure

Equations (10) and (11) are reminiscent of the QG moment recursions introduced by Barton & Turelli (1987), Turelli & Barton (1990), and Bürger (1991) for frequency-independent selection under multilocus genetics. Apart from the simpler genetic assumptions, the main technical difference with their approach is that we derive these equations at the morph level and only leading-order terms are kept under the small morph variance approximation. In effect, by allowing for the interaction between the different morphs, OMD effectively extends these moment methods to multimodal distributions, frequency-dependent selection, and a broad range of ecological scenarios.

Of course, as typical of moment methods, the system of equation is not closed. To make progress, we need to rely on a moment-closure approximation. The simplest is to assume that each morph distribution is and remains normal (so that the total distribution is a sum of Gaussian peaks). In that case, the kurtosis is 3, so that the rate of change of the morph variance is simply given by the squared variance, 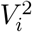 (Appendix A; Barton & Turelli, 1987; Lynch & Walsh, 1998; Sasaki & Dieckmann, 2011; Walsh & Lynch, 2018, chapter24). Note that, in contrast to the classical QG theory, the Gaussian closure is here applied at the morph level, and not at the population level. Hence, the full distribution will generically be non-Gaussian and asymmetric. Other morph-level moment closure approximations can also be used, such as the “rare alleles” or “house-of-cards” approximations (Barton & Turelli, 1987; Sasaki & Dieckmann, 2011; Walsh & Lynch, 2018, chapter 24], or beta distribution approximation (Klauschies et al., 2018; Cropp & Norbury, 2021). We briefly discuss these closure schemes in Appendix A. In Box 3, we present the full oligomorphic approximation of the resource-consumer model (Model 1 in Box 2), based on the Gaussian closure.

**Box 3: Oligomorphic dynamics of the resource-consumer model**

The joint dynamics of the density of consumers, morph frequencies, morph means, and morph variances for Model 1 in Box 2 are given by:

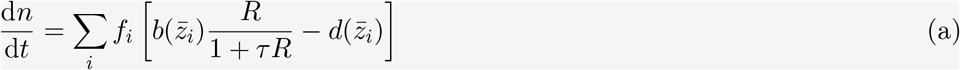

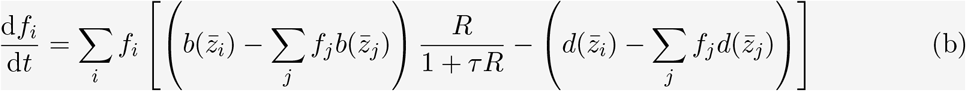

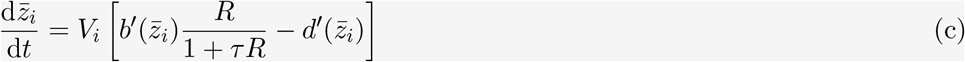

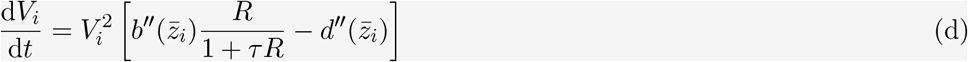

For the dynamics of morph variances, we have used a Gaussian closure approximation of the morph distribution. The dynamics of the resource can also be approximated to leading order (see Online Appendix S).

In order to clarify the connections with PG, AD, and QG, note that we recover the dynamics of single-locus PG models in the limit *V_i_* = 0 (we are then left with only equations (a) and (b)), and the dynamics of frequency-dependent QG models with only one morph and fixed variance (we are then left with equations (a) and (c)). The terms between brackets in equations (c) and (d) correspond to the first and second partial derivatives of invasion fitness in AD.

In addition, if we want to incorporate frequency-dependent competition, as in the Model 2 in Box 2, we can approximate the function *d*(*z*) = *d*_0_ + *n* ∫ *c*(*y* − *z*)*ϕ*(*y, t*)d*y* using the multi-morph decomposition and small morph variance approximation to obtain:

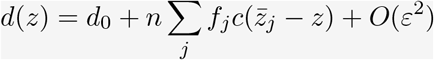

and we can plug this expression into the OMD equations to complete the analysis. Note the similarity between the latter expression and the definition given in Box 2 for a discrete phenotypic distribution. The two coincide when the morph variances are vanishingly small.

### 3.8 Separation of time scales

A very important consequence of the small morph variance approximation is that the different statistics of the model do not change on the same time scales. Typically, the population density and the morph frequencies change on a fast time scale, while the morph means and morph variances change more slowly.^1^ This means that, when genetic variation within each morph is not too large, ecological densities and morph frequencies will change quickly compared to the morph mean, and will very quickly respond to any change in the means. This is very useful as it allows for the application of mathematical quasi-equilibrium techniques (Rinaldi & Scheffer, 2000; Cortez & Ellner, 2010; Cortez & Weitz, 2014) to simplify the eco-evolutionary dynamics.

### 3.9 Comparison of the different eco-evo frameworks

Figure 1b gives an overview of the connections between the different theoretical frameworks available to study eco-evolutionary dynamics. Multi-species ecological dynamics and PG are limit cases of OMD for fixed values of the traits (e.g. *V_i_* = 0 and 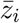 is constant for all morphs). QG correspond to the limit case of a unimodal trait distribution, typically with constant variance. The results of AD are retrieved in the limit of vanishingly small morph variances. OMD therefore acts a unifying framework allowing for a fruitful dialogue between different theoretical schools.

## 4 Applications

We now show how the OMD equations can be used to shed light on various phenomena such as the interplay between fast and slow evolution regimes, transient eco-evolutionary dynamics, disruptive selection through frequency-dependent selection, the dynamics of genetic variance under mutation and selection, and the evolution of skewed trait distributions, structured populations, and multivariate traits.

### 4.1 Bridging the gap between fast and slow evolution

OMD allows us to analyse eco-evolutionary dynamics across different relative ecological and evolutionary time scales. In particular, OMD can be used to study the role of “fast evolution” fueled by a large standing variation at the population level.

As a simple model, consider a population characterised by a bimodal distribution with two morphs. Even if the morph variances *V*_1_ and *V*_2_ are not large, the population-level variance, *V*, can be substantial if the difference in morph means, 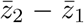 is large. As we shall see, the magnitude of the population-level variance *V* will greatly impact the form of the feedback between ecological and evolutionary dynamics.

To fix ideas, let us consider our resource-consumer example (Model 1). The mean trait, at the population level, can be calculated from the morph frequencies and morph means as 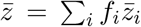. Using equations (9) and (10), we obtain

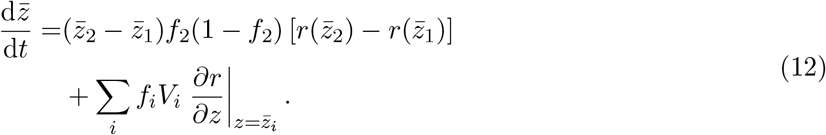

Equation (12) captures the coupling between fast and slow evolution. The first line of equation (12) represents the contribution of the dynamics of frequencies to the change in the mean (i.e. what happens when the heights of the peaks change), while the second line represents the contribution of the change in morph means (i.e. when the position of the peaks change).

When the two morphs are very different (that is, when the distance between the two peaks of the distribution is large compared to the morph variances *V*_1_ and *V*_2_), the dynamics of the mean trait in the population are dominated by the change in frequencies. Then, the change in mean trait is given by the difference in growth rates between the two morphs, scaled by the variance *f*_2_(1 − *f*_2_), as in population genetics or two-species ecological models (see e.g. equation (3)). In contrast with these approaches, however, equation (12) assumes non-zero within-morph genetic variation.

On the other hand, when the two morphs are close (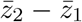 is small, so that the population can be thought of as quasi-monomorphic), the dynamics of the mean trait are dominated by the change in morph means. As in classical AD and QG approaches, the direction of selection is now given by the slope of the morph growth rates, weighted by the morph frequencies and variances.

Hence, a morph that has a higher growth rate but a lower slope could be transiently selected (on the fast time scale), but eventually counter-selected in the long run (on the slow time scale). This leads to a very distinct pattern of environmental feedback, where evolutionary dynamics (i.e. the change in the trait mean 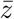) take place before the ecological dynamics have the time to settle on an ecological attractor. This is characteristic of fast evolution regimes. Once selection has eroded the population-level genetic variation, the dynamics enter the typical gradient dynamics of slow evolution regime, where gradual change in the mean trait triggers a fast readjustment of the ecological variables. Figure 2 shows that OMD can capture how different levels of initial standing variation lead to distinct patterns of eco-evolutionary feedbacks, despite identical long-term steady states. For our specific example, note that, for small initial variance, we have fast relaxation of the ecological dynamics followed by a gradual evolution of the mean trait which slowly decreases towards its evolutionarily stable value. In contrast, for large initial variance, we observe fast evolutionary dynamics characterised by a transient increase in mean trait followed by a sharp decrease, then a slow increase of the mean trait towards the ESS. These different feedbacks have distinct practical implications: with large standing variation, we expect transient selection for rapacious consumers, whereas when standing variation is small, we predict selection to always favour more prudent consumers. Similar arguments could be used to study predator-prey cycles under different fast or slow evolution, as investigated by Cortez & Weitz (2014), who studied the impact of genetic variation on the speed of evolution and the form of the eco-evolutionary feedback, using clonal models and frequency-dependent QG models. The OMD approach provides a connection between these two types of models.

**Figure 2:**
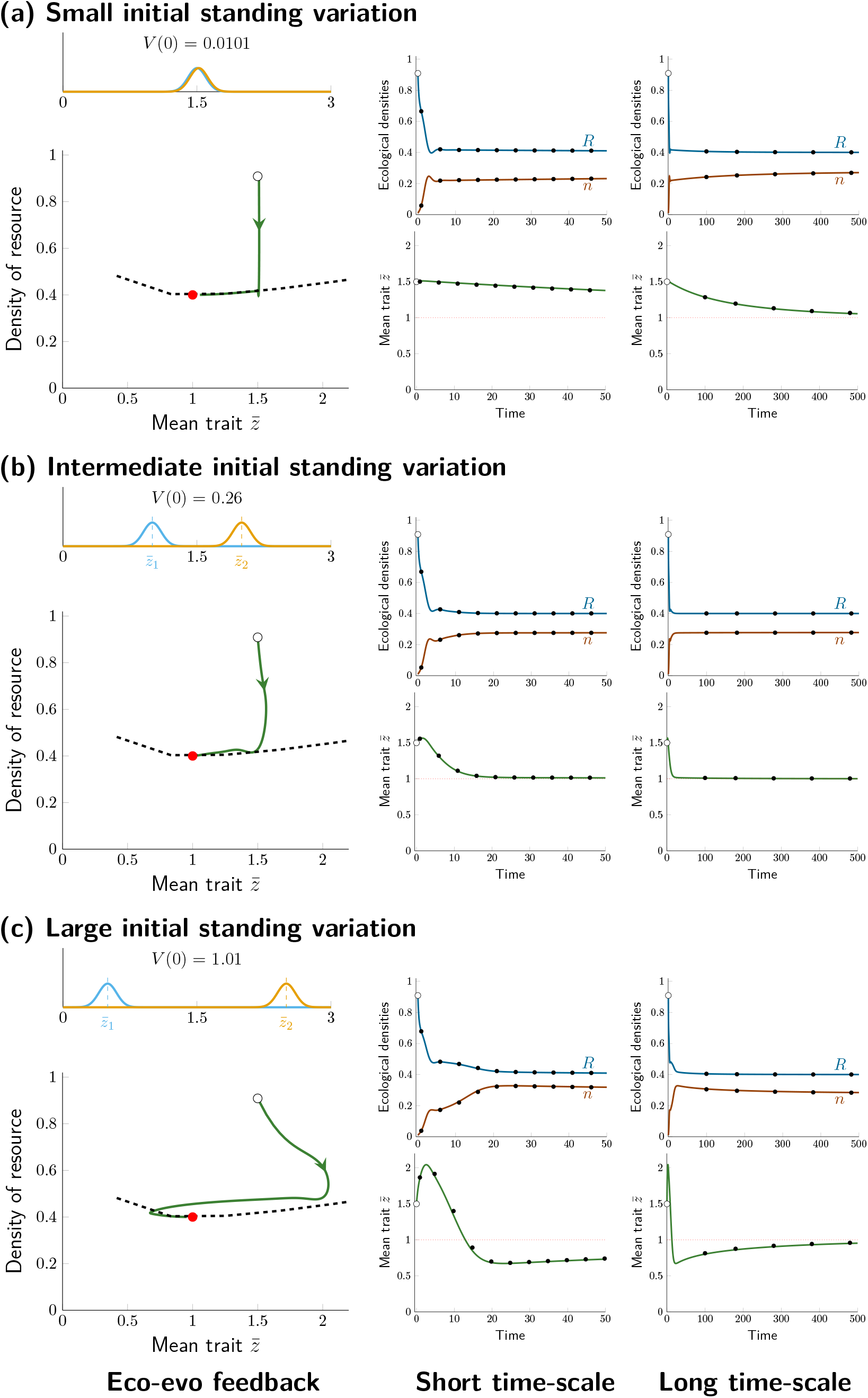
Coupling fast and slow evolution. Depending on the magnitude of the initial standing variation, the dynamics of the resource-consumer model (Model 1 in Box 2) are characterised by very distinct transient eco-evolutionary feedbacks, corresponding to different balances of fast vs. slow evolution. OMD is able to integrate the dynamics across these different time scales. The dashed line on the left panels corresponds to the quasi-equilibrium approximation for the monomorphic dynamics. In each panel, the value of the variance at the population level is given, and corresponds to the variance calculated for a population with two Gaussian morphs with morph variances *V*_1_(0) = *V*_2_(0) = 0.01 and distances between morphs 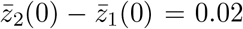 (a), 1 (b), or 2 (c). See Online Appendix S for technical details and parameter values.

Because OMD gives predictions on the morph distributions, it allows us to better understand the evolutionary mechanisms affecting the change in the trait distribution. For instance, we show in figure 3 how the two phases of fast vs. slow evolution observed in the simulations of figure 2c can be ascribed to changes in morph frequencies (peak heights) vs. morph means (peak positions). By going back and forth between the morph and population-level statistics, it is thus possible to shed light on the mechanisms shaping the dynamics of trait distributions.

**Figure 3:**
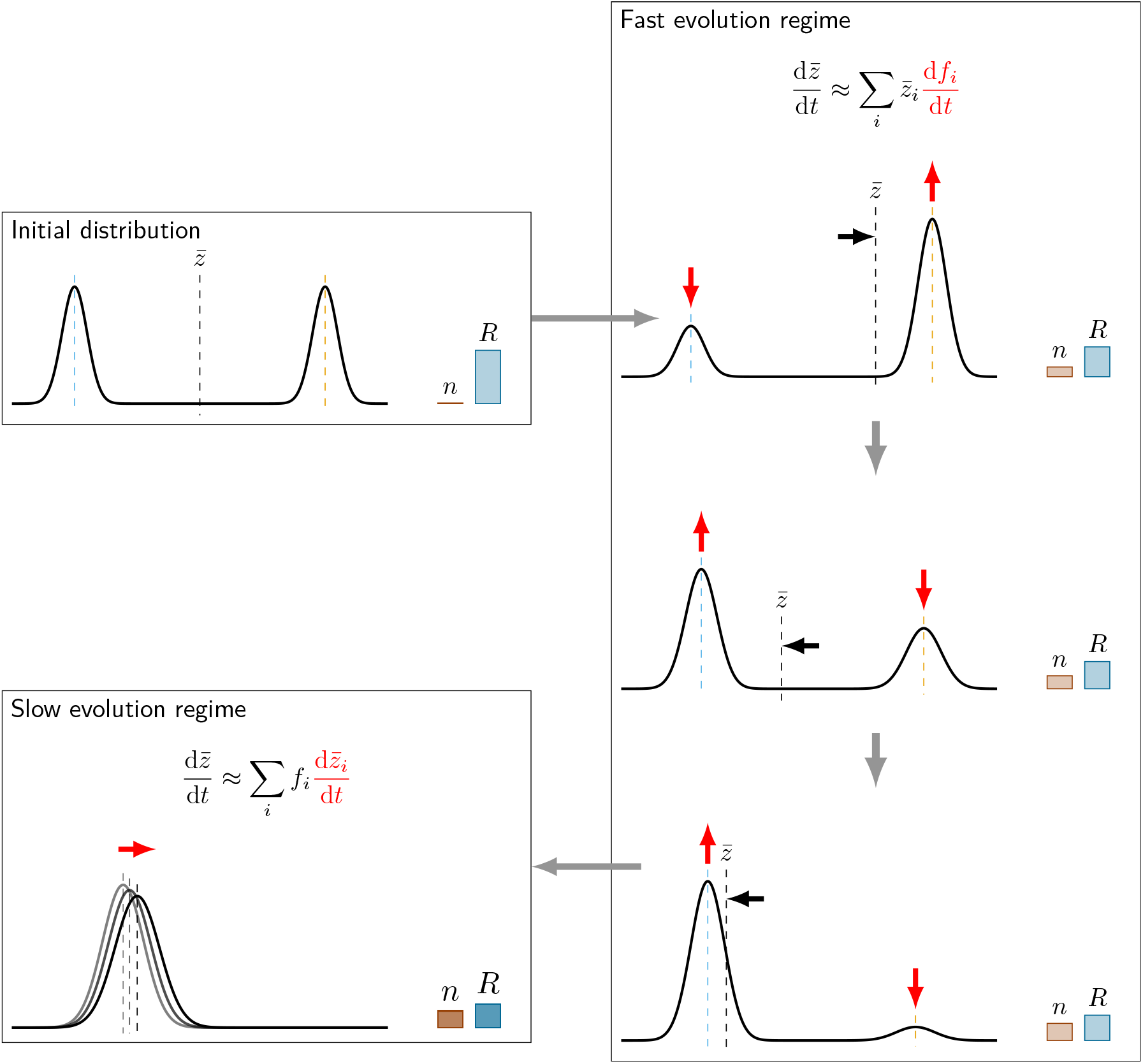
Linking fast and slow evolution to mechanisms. Starting with a bimodal distribution with a large initial standing variation at the population level (as in figure 2c), the dynamics of the resource-consumer model are first characterised by a phase of fast evolution where most of the change in the population mean is due to changes in morph frequencies while the morph means stay approximately constant. This leads to the competitive exclusion of one morph, at which stage the population enters a phase of slow evolution where the mean trait changes gradually in the direction of the selection gradient. Note that the trait variance also changes in that phase. See Online Appendix S for technical details and parameter values.

### 4.2 Transient and non-equilibrium evolutionary dynamics

An immediate consequence of the abililty of OMD to capture both fast and slow evolution is that it allows for the study of transient evolutionary dynamics, on short time scales. In principle, this makes it easier to formulate testable predictions that match the time scales of empirical observation in the field or in the lab, while retaining the ability to make long-term evolutionary predictions as in AD. We therefore think that it represents an interesting step towards a better integration of theory and data in evolutionary ecology.

In addition, because OMD does not assume that the population is always near an ecological attractor, it can be used to investigate evolution in non-equilibrium, dynamical environments. For instance, we have recently applied this approach to analyse the evolution of pathogen virulence under the repeated epidemics caused by antigenic escape (Sasaki et al., 2022). The problem of managing transient pathogen evolution over the course of an epidemic is central in evolutionary epidemiology (Lenski & May, 1994; Day & Proulx, 2004; Day & Gandon, 2007), and this could be an interesting application of OMD. For instance, if we make an analogy between consumers and pathogens on the one hand, and resources and susceptible hosts on the other hand, we can use equation (12) to study the transient evolution of more transmissible and virulent pathogen strains during an epidemic, followed by selection for more prudent pathogens once the endemic phase is reached. More generally, as the oligomorphic decomposition and small morph variance approximations do not rely on the assumption of equilibrium dynamics, we think OMD could be used to generate predictions for the evolutionary consequences of non-equilibrium processes such as seasonality, and to better understand how environmental fluctuations in space or time affect diversification.

### 4.3 Frequency-dependent disruptive selection

In contrast with QG, OMD can be used to study how disruptive selection may cause the trait distribution to split into several modes. Consider for instance a continuous-trait version of Model 2 in Box 2, where the mortality rate of individuals with trait *z* is

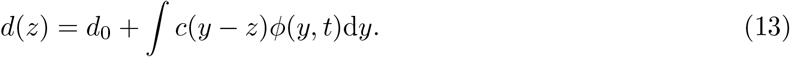

This can be used to represent the increase in mortality caused by competition between phenotypically similar individuals (Roughgarden, 1972; Slatkin, 1980). Using the oligomorphic decomposition and a double Taylor expansion (Box 3; Online Appendix S; Sasaki & Dieckmann (2011)), it is possible to approximate the change in the morph means in Model 2 as

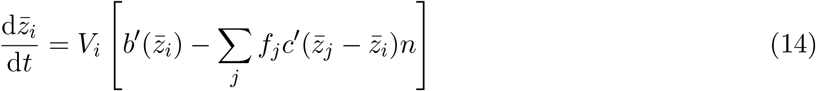

Intra-specific competition at the population level is thus approximated by the pairwise competition between the different morphs, so that the equations of the morph means are all coupled. The presence of the frequencies *f_j_* shows that, in a literal sense, equation (14) captures the notion of frequency-dependence. An analogous equation can be derived for the change in morph variance.

Using the morph dynamics, we can thus describe how, starting from a quasi-monomorphic distribution with two slightly different morphs, frequency-dependent competition leads to the two morphs moving apart until the distribution becomes bimodal. Figure 4a compares the predictions of the OMD approximation to numerical simulations of the full model without the oligomorphic decomposition, and shows that the OMD predictions capture well the timing and location of the branching although, due to the built-in morph decomposition, they tend to overestimate the speed at which the distribution splits into two modes. However, the trait distributions and population densities before and after branching are accurately predicted (figures 4b and 4c). Sasaki & Dieckmann (2011) give a detailed discussion of how the OMD predictions relate to classical character-displacement models (Roughgarden, 1972; Bulmer, 1974; Roughgarden, 1976; Slatkin, 1980; Taper & Case, 1985). The OMD approach goes one step beyond by relaxing the assumption of fixed variance (e.g. Roughgarden (1976)), Gaussian distributions (e.g. Slatkin (1980)), or single-locus two-allele genetics (Bulmer, 1974). As we have seen, all these previous models can be seen as special cases of the more general OMD framework.

**Figure 4:**
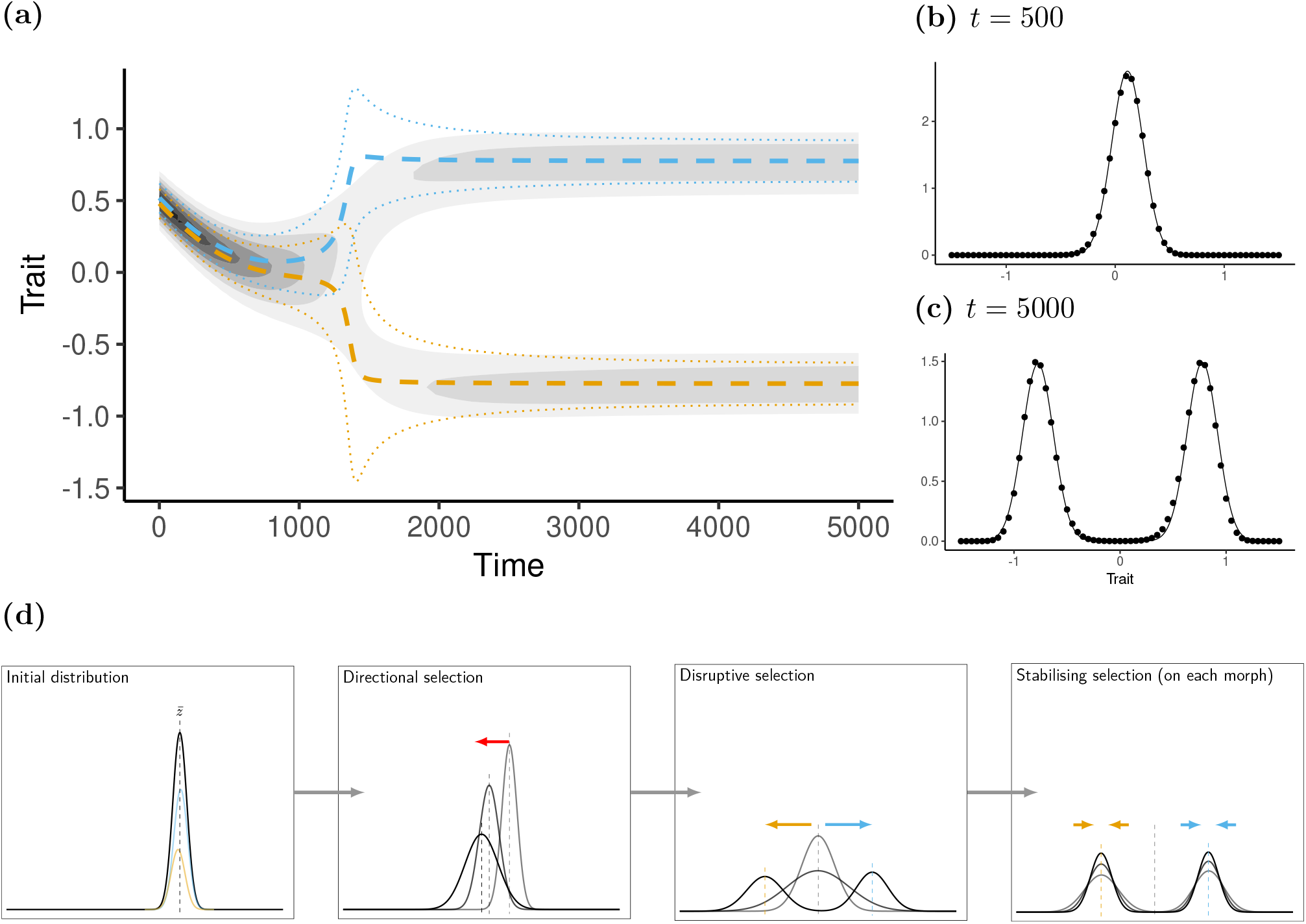
Capturing frequency-dependent disruptive selection. OMD can be used to capture how an initially unimodal distribution splits into two peaks. In panel (a), the contour plot represents the trait distribution obtained from numerical simulations of the full model (Model 2 in Box 2 without the oligomorphic approximation, see Online Appendix S). The orange and blue lines represent the results of simulations of a two-morph oligomorphic approximation, starting from initially similar morphs (dashed lines: morph means; dotted lines: morph standard deviations). Population-level trait distributions before (b) and after (c) branching are represented on the right (dots: full simulations; lines: oligomorphic approximation). In panel (d), we present, using the OMD results of panel (a), a schematic of the time-evolution of the trait distribution at the population level, *ϕ*(*z, t*). On the left panel, the initial distribution (in black) is the sum of the two morph distributions (in blue and orange). On the other panels, we show the total distribution at different time points (lighter shades of grey corresponding to earlier time points). The total trait distribution first moves towards the left under directional selection, before becoming bimodal through disruptive selection (at which point, the two morph distributions begin to move apart). Stabilising selection on each peak then leads to the equilibrium trait distribution, with each peak corresponding to a distinct morph. See Online Appendix S for technical details and parameter values.

Another question that can be addressed using OMD is to approximate the time required until an initially unimodal character distribution splits into two distinct morphs under frequency-dependent disruptive selection. For clonally reproducing species, this is a measure of the waiting time until adaptive speciation. As shown by Sasaki & Dieckmann (2011), an approximation for this waiting time can be derived under various moment closure approximations, and is found to be typically inversely proportional to the curvature of the fitness landscape at the evolutionary branching point.

### 4.4 Dynamics of genetic variance and mutation-selection balance

In contrast to the majority of QG approaches, the OMD approach allows us to capture the dynamical nature of genetic variance due to both selection and mutation. The small morph variance approximation leads to explicit expressions for the effect of selection on the dynamics of variance, in terms of the frequencies and means of the different morphs (equation (11), Sasaki & Dieckmann, 2011; Lion et al., 2022), which sheds light on the ecological processes leading to diversification. It is also possible to incorporate mutation to capture the fact that the depletion of variance due to stabilising selection will tend to be restored by the generation of genetic variation through mutation. There are various ways to incorporate mutation in OMD. The simplest is to assume unbiased mutations occurring at rate *μ* with the distribution of mutation effects having variance 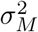 (Sasaki & Dieckmann, 2011). This leads to the following equations for the morph variances:

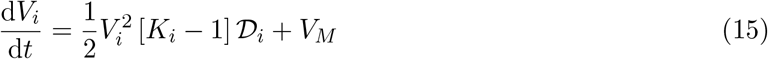

where 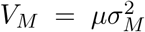 is the mutational variance and 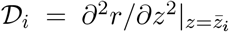 measures the curvature of the fitness landscape and therefore the strength of disruptive or stabilising selection. When 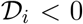, selection on morph *i* is stabilising and the morph variance equilibrates at a mutation-selection balance. Assuming that the morph distributions are Gaussian (*K_i_* = 3), the equilibrium variance can be simply calculated as

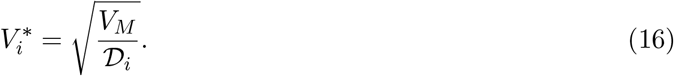

Equation (16) generalises classical expressions obtained under the assumption of Gaussian or quadratic stabilising selection (Kimura, 1965; Turelli, 1984), with the difference that the parameter measuring the intensity of selection, 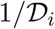, is not a constant but depends on the mean trait and on the environmental feedback. Other expressions can be obtained under other closure assumptions, such as the house-of-cards approximation (Appendix A; Turelli, 1984; Sasaki & Dieckmann, 2011).

### 4.5 Population-level variance and skewness

From the moments of the morph distributions, it is possible to calculate the moments of the population-level distribution, as we have already done for the mean in section 4.1. This allows one to investigate how the variance and skewness of the whole trait distribution change over time through selection and mutation. In particular, the fact that the skewness of the trait distribution can affect directional selection on the mean trait has been repeatedly noted in theoretical studies (Barton & Turelli, 1987; Turelli & Barton, 1990; Débarre et al., 2015), and is backed up by experimental data (Bonamour et al., 2017). However, these previous works were faced with the problem of closing the system of population-level moment equations. Because with OMD the dynamics of moments are closed at the morph level, expressions for the population-level moments can be derived that only depend on the heights, locations, and widths of the various peaks of the distribution (Box 4). From this point of view, OMD can be seen as a partial solution to the moment closure conundrum that has hampered the applicability of moment methods in QG, and has the potential to allow for a tighter integration between theoretical predictions and empirical trait distributions. Figure 5 gives a taste of how OMD can be used to track the short-term dynamics of non-Gaussian trait distributions under the action of natural selection and small mutation. Further developments will need to investigate the robustness of the moment closure approximation when mutation is stronger, but we already note that the small-morph approximation is remarkably accurate even when the morph distributions are not particularly narrow, as in figure 5.

**Figure 5:**
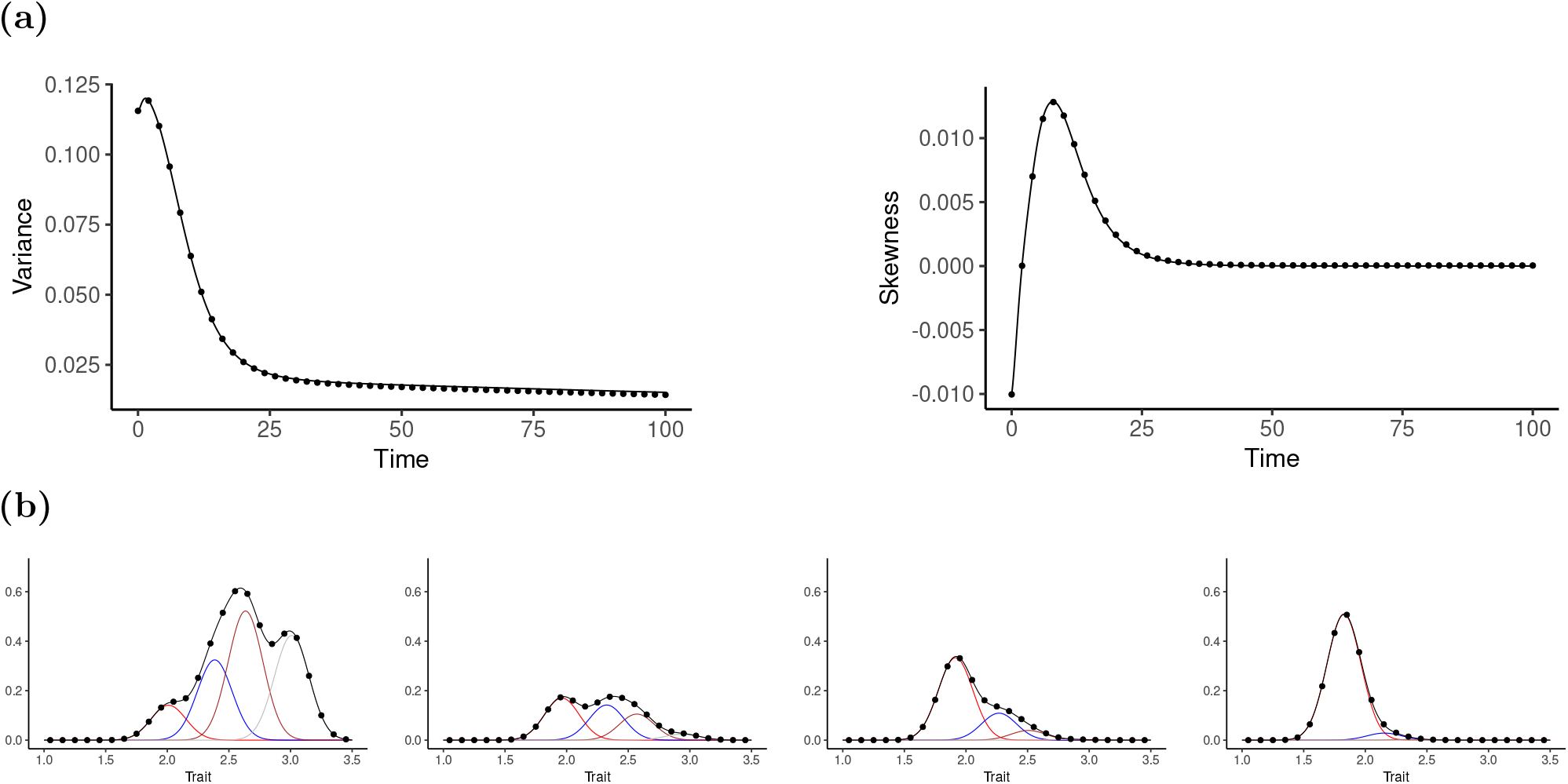
Transient dynamics of non-Gaussian trait distributions. Panel (a) shows the dynamics of the variance (left) and third central moment (right) of the trait distribution evolving in the resource-consumer model (Model 1 in Box 2). OMD (line) yields a very good approximation of the dynamics (dots are simulation results without the oligomorphic approximation). Panel (b) shows how OMD can capture the full transition from left-skewed (*t* = 0, negative skewness) to right-skewed (*t* = 10, positive skewness) to symmetric (*t* = 20) trait distribution. In both the OMD (lines) and full (dots) simulations, the trait distribution is initiated as a sum of four Gaussian peaks (coloured lines at *t* = 0). OMD tracks how these morph distributions change (coloured lines at *t* > 0), and the resulting population-level trait distribution (black lines) accurately predicts that in the full simulation (dots). See Online Appendix S for technical details and parameter values.

### 4.6 Population structure and multivariate traits

As in AD and QG, OMD can be extended to take into account class structure (Lion et al., 2022; Wickman et al., 2022) and the joint evolution of multiple quantitative traits (Sasaki et al., 2022).

Class structure is a major feature of natural biological populations, taking into account individual differences in state including age, spatial location, infection or physiological status, and species. By deriving equations for the frequencies and moments of the morph distributions in each class, it is possible to apply OMD to a broad range of realistic ecological scenarios (Lion et al., 2022; Wickman et al., 2022). Furthermore, Lion et al. (2022) show how the theory of reproductive values can be used to obtain compact analytical expressions for the dynamics of multimodal trait distributions in structured populations under density- and frequency-dependent selection. From a biological perspective, this sheds light on how the quality and quantity of individuals in different classes affect the eco-evolutionary dynamics.

**Box 4: Calculating population-level moments from morph moments**

The mean of a multi-morph distribution can be easily calculated as a function of the morph means and frequencies. This yields the following intuitive expression:

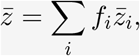

which we can use to derive an equation for the dynamics of the population-level mean (see e.g. section 4.1).

For the variance and third central moment of the trait distribution, similar, albeit more complex, expressions can be derived. For the variance, we have:

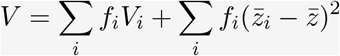

The first term is the average of the morph variances, weighted by morph frequencies. The second term is the weighted average of the squared deviations of the morph means from the population-level mean. This captures the fact that the variance of the trait distribution increases either when the morph distributions are flatter or when the morphs move apart. This relationship makes it possible to track the dynamics of the population variance from those of the morph moments.

The third central moment, *T*, of the population-level distribution can similarly be expressed as a function of the morph frequencies, means, and variances. For two morphs, this can be written as

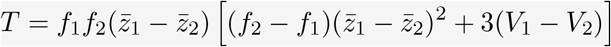

under the assumption that each morph distribution is symmetric. In that case, the above expression captures the fact that a bimodal distribution will be skewed if its two peaks have different heights or widths.

The analysis of multivariate quantitative traits is a key feature of QG models. With OMD, it is possible to obtain dynamical equations for the change in morph means and variances (Box 5; Sasaki et al. (2022)) that closely resemble those derived in QG by assuming Gaussian unimodal distributions (Lande & Arnold, 1983; Phillips & Arnold, 1989; Mullon & Lehmann, 2019). However, OMD goes one step further by allowing us to take into account selection on multimodal and skewed trait distributions, as well as frequency-dependent disruptive selection caused by environmental feedbacks. There are interesting connections with QG community ecology models (Barabás & D’Andrea, 2016; Barabás et al., 2022), which retain the classical QG assumption of trait normality but allow for multi-species ecological interactions, and yield equations that are very similar to those describing the morph dynamics in Box 5.

## 5 Perspectives

There are many directions in which to take OMD, but the main theoretical challenges, from a biological point of view, are to take into account complex genetical systems, environmental variance, demographic and environmental stochasticity, and integration with data. We discuss these in turn.

### 5.1 Genetics, inheritance, and mutation

Most of the theory we describe here has been developed under the assumption of clonal reproduction and therefore leaves aside the potential complexities of genetic architecture (e.g. ploidy, number of loci, interaction across loci, and sexual reproduction). This is a classical simplification shared by most PG, AD, and QG models interested in the interplay between ecology and evolution. In fact, although the mathematical justification of QG often invokes random mating in sexually reproducing organisms, most eco-evolutionary QG models do not explicitly model sexual reproduction and rely on equations that are identical to those derived for clonally reproducing organisms. There are however good empirical and theoretical reasons for arguing that genetic architecture and variation may have strong effects on eco-evolutionary processes, especially on the short term (Barrett & Schluter, 2008; Bolnick et al., 2011; Yamamichi, 2022), and on the shape of trait distributions (Yeaman & Guillaume, 2009; Débarre et al., 2015). It is also known that sexual reproduction affects disruptive selection and that assortative mating is often necessary for diversification to occur in sexually reproducing organisms (Doebeli et al., 2007). Coupling OMD with more complex genetic models will therefore be an important challenge for future extensions of the theory. Although the moment equations derived in QG models assuming sexual reproduction have been found to be similar to those derived under clonal reproduction (Bürger, 1991; Débarre et al., 2013; Barabás & D’Andrea, 2016; Barabás et al., 2022), additional assumptions are often needed when dealing with multi-locus genetics and recombination (see e.g. the discussion in Débarre et al. (2013)). At a technical level, further developments in this area may be fostered by analysing the connections between OMD, classical multi-locus moment equations (Barton & Turelli, 1987; Bürger, 1991; Turelli & Barton, 1994), and recent theoretical developments that have studied how sexual reproduction and genetic architecture affect the origin and maintenance of polymorphism (e.g. Patel & Bürger (2019) and Dekens et al. (2021)). Biologically, these technical extensions could be very fruitful for our understanding of the complex feedback between ecological dynamics and genetics.

**Box 5: Joint evolution of multiple quantitative traits**

Following the seminal work of Lande & Arnold (1983), the dynamics of multivariate quantitative traits in QG models are given by the product of a genetic (cov)variance matrix **G** and of a selection gradient. As shown in Sasaki et al. (2022), the multivariate extension of OMD takes a similar form at the morph level:

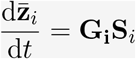

where **G**_*i*_ is the morph-specific genetic (co)variance matrix and **S**_*i*_ is the gradient of the fitness function *r*(**z**, **E**) with respect to the multivariate trait **z**.

As in the univariate case, the dynamics of the morph means depend on a dynamical measure of genetic variation, **G**_*i*_, which changes as follows

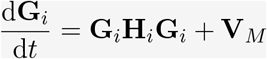

where **H**_*i*_ is the Hessian of the fitness function for morph *i* and **V**_*M*_ is the diagonal matrix of the mutational variances for each trait. Similar equations have been derived in multi-species models (Débarre et al., 2014) or single-species models with a Gaussian distribution (Mullon & Lehmann, 2019).

In addition, we have assumed, for simplicity, a one-to-one relationship between the genotype and phenotype, and ignored the possibility that individual or environmental variation may alter the phenotypic value for a fixed genotypic value. As a result, there is no distinction between the additive genetic variance and the phenotypic variance in our morph-specific moment equations (in other words, heritability is assumed to be 1 throughout this paper). Of course, this simplification contrasts with the large and successful literature in theoretical and statistical QG, which is specifically devoted to taking into account how the phenotypic distribution is shaped by both genetical and environmental effects (Falconer, 1960; Lynch & Walsh, 1998; Kruuk et al., 2018). However, it would be relatively straightforward to extend OMD in that direction, by writing the phenotypic distribution as a joint distribution of genotypic and environmental effects, and integrating over the environmental distribution (see e.g. the Appendix of Iwasa et al. (1991)). More generally, it would be particularly interesting to combine the oligomorphic decomposition and small morph variance approximation with well-established statistical QG methods. This could be done by building on existing multi-species QG theory (e.g. Barabás & D’Andrea (2016) and Barabás et al. (2022)), or by coupling OMD with the framework of Coulson et al. (2017) which tracks both the genetical and environmental components of phenotypic distributions.

One of the main limitation of OMD so far is that the accuracy of the approximation can be eroded in the limit of strong mutation, as this may cause the small morph variance approximation to break down. To solve this, inspiration could come from other related approaches that have been developed to handle high mutation rates on the stationary trait distributions, but lack the dynamical aspect of OMD (Mirrahimi & Gandon, 2020). There are also interesting theoretical challenges in better understanding how various mutation models affect the robustness of OMD.

### 5.2 Population and community structure

Non-genetic heterogeneity between individuals is a key feature of natural populations, which is known to have important ecological and evolutionary consequences (Coulson & Tuljapurkar, 2008; Ozgul et al., 2009). In demography or ecology, this heterogeneity is taken into account by adding a discrete or continuous structure to the population (Charlesworth, 1994; Caswell, 2001). In addition to the class-structure extension of OMD we discussed above, it would be interesting to apply the oligomorphic decomposition and small morph variance approximation to simplify the dynamics of eco-evolutionary integral projection models (IPMs), which have been developed to model the dynamics of the distribution of heritable traits, typically under QG assumptions, and successfully applied to analyse empirical data (Barfield et al., 2011; Childs et al., 2016; Rees & Ellner, 2016; Coulson et al., 2017). OMD could be used to decompose the population-level trait distribution and translate the IPM into morph-specific equations. As we have shown, OMD allows for a decomposition of fast and slow evolution regimes and for the study of multimodal trait distributions and complex environmental feedbacks, which would be an interesting contribution to current IPM approaches.

Another interesting extension would be to use OMD to better understand eco-evolutionary dynamics at the community level. Over the past 20 years or so, there has been a renewed interest in incoporating evolutionary considerations into community ecology (Jansen & Mulder, 1999; Norberg et al., 2001; Fussmann et al., 2007; Johnson & Stinchcombe, 2007; Norberg et al., 2012; Ellner, 2013; Weber et al., 2017; Nordbotten et al., 2020; Govaert et al., 2021). Most models rely on the assumptions of PG or QG to introduce genetic variation into community dynamics, and resemble the oligomorphic equations if we equate one morph to one species (Norberg et al., 2001; Klauschies et al., 2018). But OMD could be used to allow for multiple morphs within each species and to analyse how frequency-dependent selection, multimodal or skewed trait distributions, and rapid evolutionary processes affect community stability and evolution.

### 5.3 Demographic and environmental stochasticity

Although demography and evolution are inherently stochastic processes, many existing developments of OMD focus on deterministic models, instead of formulating a stochastic model as in traditional population genetics approaches. Thus, in its current form, OMD relies on a large-population assumption, like most AD models (Rousset, 2004; Champagnat et al., 2006; Méléard, 2011; Lehmann et al., 2016). The benefit of this simplification is that it makes the model more tractable to analyse the coupled eco-evolutionary dynamics without resorting to the assumptions of frequency- or density-independence often encountered in population genetics (Holt & Gomulkiewicz, 1997; Heino et al., 1998; Rice, 2004; Day, 2005). The drawback is that a deterministic framework cannot capture the fact that both ecological (Lande et al., 2003) and evolutionary (Lenormand et al., 2009) processes can be strongly affected by demographic or environmental stochasticity.

There are however exceptions to the focalisation on deterministic models in OMD. For instance, Débarre & Otto (2016) have shown that it is possible to use the small variance approximation to analyse eco-evolutionary dynamics under demographic stochasticity. This analysis paves the way for a multi-morph extension that could be used to incorporate demographic stochasticity and multi-morph eco-evolutionary dyamics. In addition, it would be particularly interesting to further extend the OMD framework to account for environmental stochasticity, which is a key feature of many QG models (Chevin et al., 2017). Understanding the joint effect of stochastic population dynamics and environmental feedback on eco-evolutionary dynamics is a very active and promising area of research in mathematical biology (Champagnat et al., 2022).

### 5.4 How many morphs?

So far, we have left open the question of how many morphs we need to consider. Mathematically speaking, the number of morphs is arbitrary, as it is possible to consider morphs with negligible frequencies or nearly equal means. However, we know that there is potentially a limit to the effective number of morphs that can be supported by the ecological dynamics under study. In addition, OMD does not allow for the splitting of a morph distribution into two, so if we initiate the dynamics with only one morph but evolutionary branching takes place, we won’t observe diversification but rather that the variance of the single morph blows up (Sasaki & Dieckmann, 2011). Hence, the number of morphs used to decompose the trait distribution will in general depend on the question we are interested in. However, the upper bound for the number of coexisting morphs will correspond to the dimension of the environment, as defined in AD, that is the effective number of ecological variables that are controlled by the population dynamics and needed to describe the effect of the environment on the sign of invasion fitness (Mylius & Diekmann, 1995; Metz et al., 2008; Metz & Geritz, 2016; Lion & Metz, 2018). Hence, existing theory can be used to guide our choice of the number of morphs. In simple cases, intuition can be used. For instance, in a two-habitat or two-resource model in a constant environment, it makes sense to start with a two-morph decomposition.

### 5.5 Application to data

A key limitation to the application of AD is that there is a separation between the theoretical outcomes and empirical data. OMD retains the ecological grounding of AD but moves beyond forecasting the long-term evolutionary stable states, and yields predictions on the dynamics of quantities that can be measured on the time scale of an empirical or experimental study, such as the mean phenotype in the population, or the heights of the different peaks of a multimodal distributions. Predictions on the shape and dynamics of the full trait distribution under standing variation can also be obtained, which allows for a tighter integration with empirical trait distributions. As a result, we think coupling OMD, empirical quantitative genetics, and community ecology would represent a fruitful research programme with the potential to lead to a much better understanding of trait and population dynamics in nature. For instance, within the large empirical literature on rapid evolutionary processes (Thompson, 1998; Hairston et al., 2005; Post & Palkovacs, 2009; Ellner, 2013; Hendry, 2017; De Meester et al., 2019), a major question is to quantify how much of the change in an ecologically relevant variable is due to evolutionary change (due to genetic or non-genetic causes) or to ecological change (Hairston et al., 2005; Ellner et al., 2011). OMD could potentially be used to shed light on how multimodality and the dynamics of variance may affect the relative magnitude of these two terms. Another example of integration with empirical data would come from applying OMD to long-term studies of wild populations often analysed using integral projection models (Ozgul et al., 2010; Coulson et al., 2017; Simmonds et al., 2020), in order to take into account skewed trait distributions (Bonamour et al., 2017) or disruptive selection. Robust statistical methods to decompose a multimodal distributions into a mixture of unimodal distributions are available (McLachlan & Peel, 2000) and already used (although not commonly) in functional ecology (Laughlin et al., 2014) and quantitative genetics (Lynch & Walsh, 1998; Gianola et al., 2006).

## 6 Conclusions

To sum up, we propose that oligomorphic dynamics will allow for a better understanding of the role of ecological feedbacks, frequency- and density-dependent selection in nature, and has the potential to facilitate a tighter integration between eco-evolutionary theory and empirical data. It is interesting to note that both the oligomorphic decomposition and the small morph variance approximation have been independently introduced in various fields over the last thirty years. For instance, they form the backbone of trait-based multi-species models in marine ecology (Wirtz & Eckhardt, 1996; Merico et al., 2014) and community ecology (Norberg et al., 2001, 2012; Klauschies et al., 2016, 2018; Nordbotten et al., 2020; Cropp & Norbury, 2021). In this perspective, we have chosen to build on the formalism introduced by Sasaki & Dieckmann (2011), which focuses on morph frequencies instead of species abundances and is thus conceptually better suited to take into account fast evolution due to rapid changes in morph frequencies. However, a major objective for future research would be to tighten the integration between this framework, community ecology theory, and ecological QG methods. Although we have focused on relatively simple ecological models for didactical puroposes, we also think that a stimulating research direction, at the interface between theory and experiments, would be to use OMD to investigate how complex ecological dynamics, such as predator-prey cycles or non-equilibrium dynamics, can be affected by relatively rapid evolutionary processes, and how genetic variation impacts the speed and outcome of population dynamical processes, notably extinction risk and evolutionary rescue (Yoshida et al., 2007; Bolnick et al., 2011; Gonzalez et al., 2013). On the technical side, there is a deep interest in the rigorous mathematical justifications of the oligomorphic decomposition and small variance approximation in the mathematical community (see e.g. Mirrahimi & Gandon (2020), Dekens et al. (2021), and Champagnat et al. (2022)), which paves the way for further theoretical developments.

Hence, there is a large body of theory on which to build to further develop OMD, with the triple ambition of working towards an improved theoretical synthesis between ecology and evolution, a better integration between theory, experiments, and empirical studies, and a better communication between theoretical schools. At a technical level, OMD moves the field on by relaxing the assumptions of unimodal trait distributions (typical of QG models), and of negligible within-morph variance and rare mutations (used in AD). At a biological level, we think OMD can be used to better understand the feedback between ecological processes and the evolution and maintenance of diversity, as well as the interplay between ecological and evolutionary dynamics across potentially overlapping time scales.

## Acknowledgements

We thank three anonymous reviewers for very helpful and insightful comments and suggestions. This study was supported by ANR JCJC grant ANR-16-CE35-0012-01 to SL, grants NIH/R01 GM122061-03 and NSF-DEB-2011109 to MB, MEXT grant JP-24115001 to AS, and the ESB Cooperation Program, The Graduate University for Advanced Studies, SOKENDAI. This publication is funded in part by the Gordon and Betty Moore Foundation through grant GBMF10578 to MB. This work was initiated during a visit by SL and MB to SOKENDAI in January 2019. SL also wishes to acknowledge Duthie, L. Govaert, V. Luque, K. Lyberger and S. Patel for stimulating discussions while writing the first draft of the manuscript.

## Appendix A Deriving OMD equations: an overview

In this appendix, we give the key steps used to derive OMD equations. Additional technical details can be found in Sasaki & Dieckmann (2011), Lion et al. (2022), and Sasaki et al. (2022). For simplicity, we drop any explicit dependency on time and environmental feedback, so we will use derivatives instead of partial derivatives with respect to time.

## Dynamics of morph frequencies and distributions

To derive the dynamics of morph frequencies and distributions, we assume that the per-capita growth rate of an individual with phenotype *z* is the same irrespective of the morph they belong to. This means that, for any morph *i*, the density *n_i_*(*z, t*) of individuals with phenotype *z* belonging to morph *i* grows at the same rate as the total density of individuals with phenotype *z*, *n*(*z, t*). Noting that *f_i_*(*t*)*ϕ_i_*(*z, t*) = *n_i_*(*z, t*)/*n*(*t*) and *ϕ*(*z, t*) = *n*(*z, t*)/*n*(*t*), this leads to the following equalities

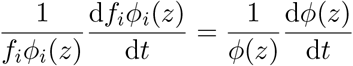

which can be rewritten as

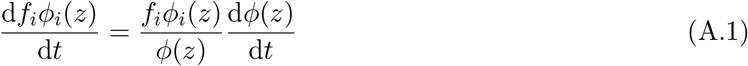

where

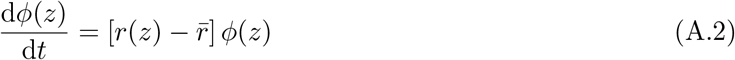

with 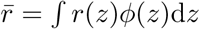 the mean growth rate of the population. Using equation (A.2) and integrating equation (A.1) over *z* then directly leads to the dynamics of the morph frequencies *f_i_*

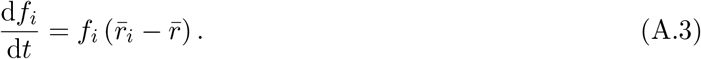

where 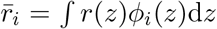 is the mean growth rate of morph *i*. To obtain the dynamics of the morph distribution, we rearrange equation (A.1) as

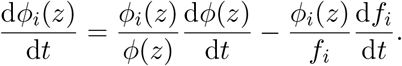

Plugging in equations (A.2) and (A.3), we finally obtain

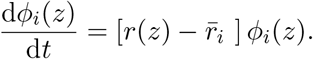

## Dynamics of morph moments

We can then calculate the dynamics of the mean and variance of the morph distribution by multiplying by *z* and 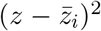 respectively and integrating over *z*. This yields

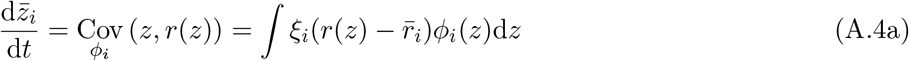

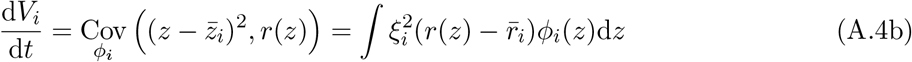

where the covariances are taken over the morph distributions *ϕ_i_*(*z*) and 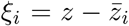 is the deviation from the morph mean.

To make further progress, we approximate the covariances in equations (A.4) with a small morph variance approximation which is obtained by Taylor-expanding the fitness function near the morph means:

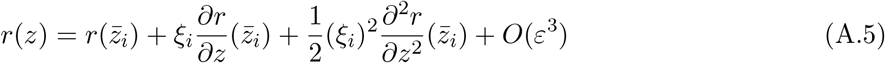

where 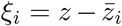 is small and *O*(*ε*). Integrating over the morph distribution *ϕ_i_*(*z*) yields the following approximation for the morph mean growth rate:

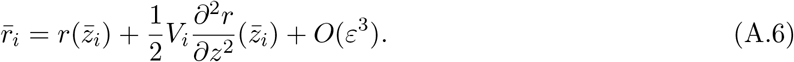

Equations (A.5) and (A.6) are then plugged into equations (A.3) and (A.4) to derive equations (9), (10), and (11). Note that, in doing so, we use the additional assumption that the morph distributions are symmetrical, so that the third central moments 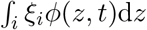 can be set to zero. The dynamics of the total density (equation (8)) can also be calculated by noting that 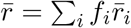 and using equation (A.6).

## Moment closure

The dynamics of morph variance are given by

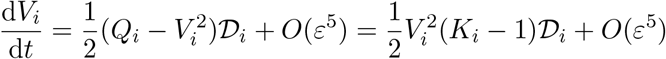

where 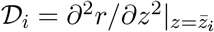 is the curvature of the fitness function at the morph mean, 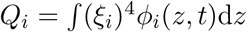 is the fourth central moment of the morph distribution, and 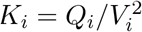 is the kurtosis. Two common moment closure approximations can then be used to close the system. The first is to assume that the morph distribution is Gaussian, so that 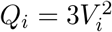, which implies *K_i_* = 1 and leads to

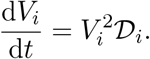

At a genetic level, this assumption can be justified by a mutation model with small Gaussian increments (see e.g. Lande, 1975; Turelli, 1984; Barton & Turelli, 1987; Walsh & Lynch, 2018, chapter 24).

A second approximation is related to the “rare alleles model” (Barton & Turelli, 1987) or “house-of-cards approximation” (Turelli, 1984), which both assume that selection is stronger than mutation and leads to a proportionality relationship between the fourth and second moments, *Q_i_* = *θV_i_* (Turelli, 1984; Barton & Turelli, 1987; Walsh & Lynch, 2018, chapter 24). Then, neglecting the 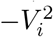 term leads to (Sasaki & Dieckmann, 2011)

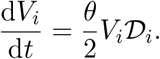

If *V_i_* is small, its dynamics are thus expected to be faster under this approximation than under the Gaussian closure (Barton & Turelli, 1987).

Finally, we note that other authors have also proposed a beta distribution approximation (Klauschies et al., 2018; Cropp & Norbury, 2021) that could also be particularly interesting to take into account the possibility of skewed morph distributions.

## Appendix S: Online Appendix

In this Online Appendix, we give the technical details and parameter values allowing potentially interested readers to reproduce figures 2–5. We also provide a succint derivation of the discrete-time OMD equations.

## Figure 2: Coupling fast-slow evolution

The figures are obtained by numerically integrating the oligomorphic dynamics of the resource-consumer model (Box 3 in the main text) together with the dynamics of the resource:

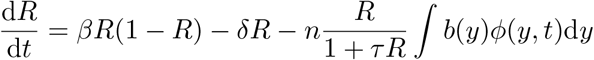

Using the oligomorphic decomposition and small morph variance approximation, the integral in the last term can be approximated as follows

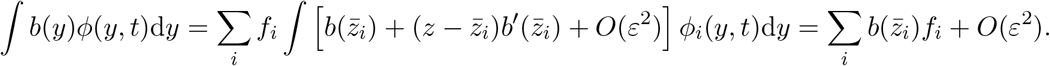

Furthermore, we use the following functional forms:

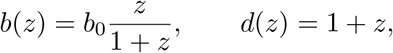

and parameter values:

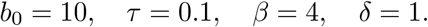

We also assume that the mutational variance is *V_M_* = 10^−5^. Initial conditions for the simulations are as follows:

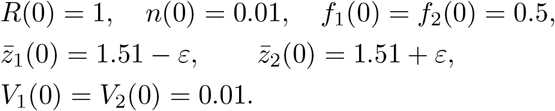

Depending on the value of *ε*, we have different levels of initial standing variation. We use *ε* = 0.01 (panel a), *ε* = 0.5 (panel b), and *ε* = 1 (panel c). The OMD predictions are compared to numerical simulations of the full model, without the oligomorphic approximation (see companion Mathematica notebook `LionEtal-OMD-SOM-Figure2.nb` for details).

On the left panels of figure 2, we plot the density of resource *R*/(1 + *τ R*) as a function of the mean trait at the population level, 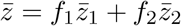. The dashed line represents the quasi-equilibrium approximation of the resource in a monomorphic population. That is, with only one morph and a narrow morph distribution, the dynamics of *n* and *R* are much faster than the dynamics of 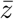, and we obtain the quasi-equilibrium expression of *R* by setting the right-hand side of the dynamics of *n* to zero. This gives

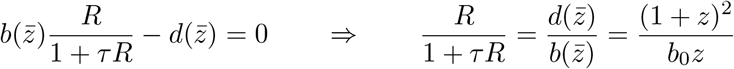

which corresponds to the dashed line in the left panels of figure 2.

The simulations of the oligomorphic approximation are compared to the numerical simulations of the full model, which are obtained by integrating the following reaction-diffusion model over a discretised trait space:

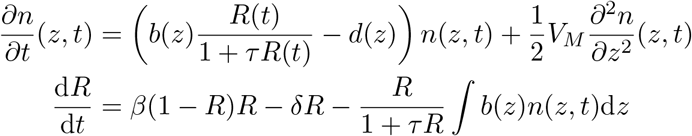

See companion Mathematica notebook `LionEtal-OMD-SOM-Figure2.nb` for additional details.

## Figure 3: Linking fast and slow evolution to mechanisms

This is meant to be a diagram, but we use real simulations to produce the figure. We use the same parameter values and initial conditions as in figure 2, except for the mutational variance which is set at *V_M_* = 10^−3^. See companion Mathematica notebook `LionEtal-OMD-SOM-Figure2.nb` for details on the code.

## Figure 4: Capturing frequency-dependent disruptive selection

To analyse this question, we turn to Model 2 in Box 2. This can be obtained from Model 1 by removing the explicit dynamics of the resource (so we always fix *R* = 1 and *τ* = 0) and assuming that competition between individuals increases mortality.

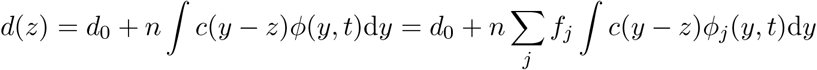

Following Sasaki & Dieckmann (2011), we Taylor-expand *c*(*y* − *z*) around 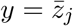 and obtain

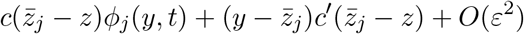

After integration over *y*, we obtain

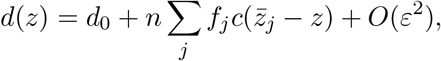

which is the approximation we use in our OMD equations. We further assume the following functional forms:

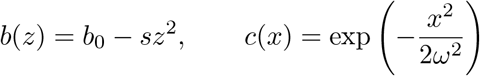

with the parameters *b*_0_ = 1, *d*_0_ = 0, *s* = 0.1, *ω* = 2, and mutational variance *V_M_* = 10^−5^. We integrate the OMD equations of Box 3 with 2 morphs, starting from initial conditions:

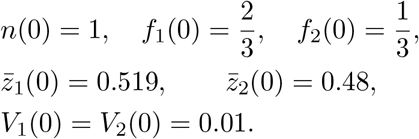

The OMD predictions are compared to numerical simulations of the full model, without the oligomorphic approximation (see companion Mathematica notebook and R script for details).

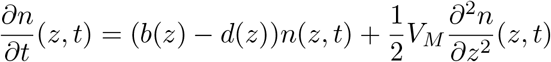

with

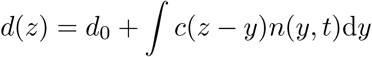

See the companion Mathematica notebook `LionEtal-OMD-SOM-Figure4.nb` for additional details.

## Figure 5: Transient dynamics of trait distributions

We use the same resource-consumer model and parameter values as in figure 2, with the following differences:

- We use *M* = 4 morphs.
- The mutational variance is *V_M_* = 10^−4^.
- The initial conditions are:

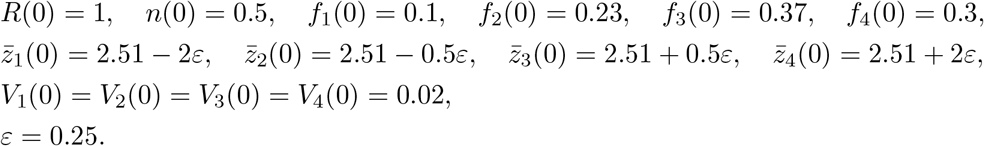

The simulations of the full model (without the oligomorphic approximation) are performed as for figure 3. See the companion Mathematica notebook `LionEtal-OMD-SOM-Figure5.nb` and the corresponding R script for details on the code.

## Discrete-time oligomorphic dynamics

Let *w*(*z, t*) be the fitness function at time *t* for phenotype *z*. During an interval Δ*t*, the trait distribution changes as

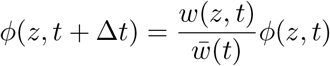

where 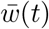 is the mean fitness. The oligomorphic decomposition leads to

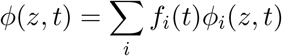

and we assume (as in Appendix A) that a phenotype *z* in morph *i* grows at the same rate as in the total population, thus

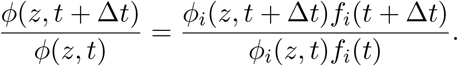

With some rearrangements, we obtain

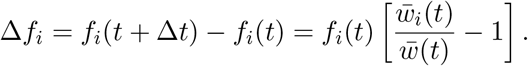

With the small morph variance approximation, we obtain

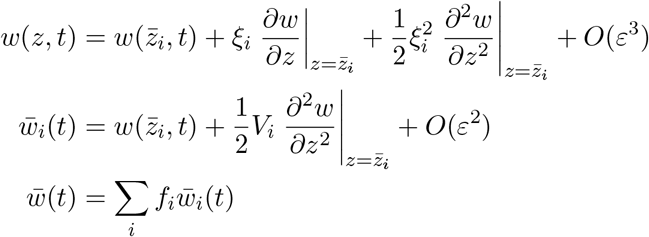

and it is easy to deduce

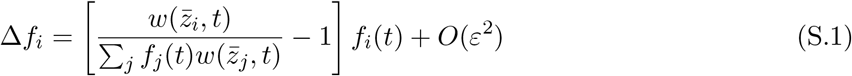

and for the total density *n*(*t*)

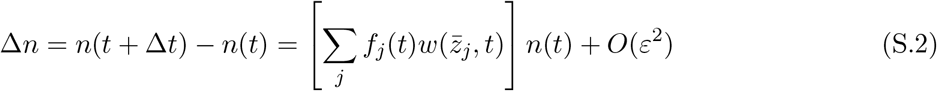

For the mean of the morph distributions, we proceed as in Appendix A and obtain

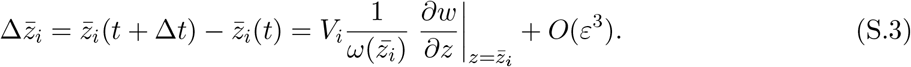

For the variance of the morph distributions, there is a subtlety, and we obtain after some calculations

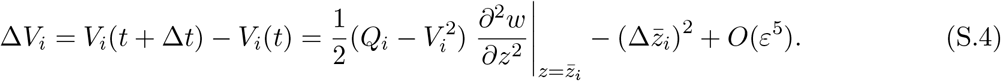

Hence, we have an extra term compared to equation (11), which corresponds to the square of the selection gradient. This is similar to classical expressions for the dynamics of the genetic (co)variance matrix **G** in QG models (Lande & Arnold, 1983; Mullon & Lehmann, 2019).

Importantly, in the continuous-time model (Appendix A), this term is absent. One way to see why is to write

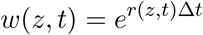

so that for small Δ*t* we have

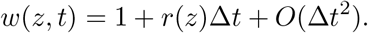

Thus, we can rewrite the dynamics of the first and second morph moments as

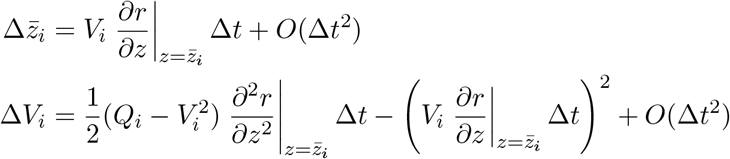

Dividing by Δ*t* and taking the limit Δ*t* → 0, we recover the continous-time equations (10) and (11). The key step here is that the square of the selection gradient is *O*(Δ*t*^2^) and therefore negligible in the continuous-time limit.

All the other OMD equations remain similar between the continuous-time and discrete-time formalisms, if we replace the growth rate *r*(*z*) by the relative fitness 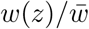.

## Footnotes

1 More precisely, under the Gaussian moment closure, the rate of change of the ecological densities and morph frequencies is *O*(1), while it is *O*(*ε*^2^) for morph means and *O*(*ε*^4^) for morph variances.

